# Mechanisms associated with pyrethroid resistance in populations of *Aedes aegypti* (Diptera: Culicidae) from the Caribbean coast of Colombia

**DOI:** 10.1101/2020.01.23.916577

**Authors:** Paula X. Pareja-Loaiza, Liliana Santacoloma Varon, Gabriela Rey Vega, Doris Gómez-Camargo, Ronald Maestre-Serrano, Audrey Lenhart

## Abstract

*Aedes aegypti* is the main vector of dengue, chikungunya, and Zika viruses, which are of great public health importance in Colombia. *Aedes* control strategies in Colombia rely heavily on the use of organophosphate and pyrethroid insecticides, providing constant selection pressure and the emergence of resistant populations. In recent years, insecticide use has increased due to the increased incidence of dengue and recent introductions of chikungunya and Zika. In the present study, pyrethroid resistance was studied across six populations of *A. aegypti* from the Caribbean coast of Colombia. Susceptibility to λ-cyhalothrin, deltamethrin, and permethrin was assessed, and resistance intensity was determined. Activity levels of enzymes associated with resistance were measured, and the frequencies of three *kdr* alleles (V1016I, F1534C, V410L) were calculated. Results showed variations in pyrethroid susceptibility across *A. aegypti* populations and altered enzyme activity levels were detected. The *kdr* alleles were detected in all populations, with high variations in frequencies: V1016I (frequency ranging from 0.15–0.70), F1534C (range 0.94–1.00), and V410L (range 0.05–0.72). In assays of phenotyped individuals, associations were observed between the presence of V1016I, F1534C, and V410L alleles and resistance to the evaluated pyrethroids, as well as between the VI_1016_/CC_1534_/VL_410_ tri-locus genotype and λ-cyhalothrin and permethrin resistance. The results of the present study contribute to the knowledge of the mechanisms underlying the resistance to key pyrethroids used to control *A. aegypti* along the Caribbean coast of Colombia.

## Introduction

*Aedes aegypti (Stegomyia aegypti)* (Linnaeus, 1762) is the main vector of the dengue (DENV), chikungunya (CHIKV), and Zika (ZIKV) viruses. The diseases caused by these viruses are of growing public health importance worldwide owing to increased proliferation of mosquito populations, increased urbanization, as well as climatic and other environmental conditions suitable for transmission [1].

Globally, the burden of disease caused by dengue is increasing; it is estimated that approximately 390 million dengue infections occur each year, of which 96 million manifest clinically [2]. In 2015, 2.35 million cases of dengue were reported in the Americas, of which >10,200 cases were diagnosed as severe dengue, causing 1,181 deaths. In Colombia, dengue is considered a public health priority owing to its endemic transmission as well as the increased occurrence of severe dengue outbreaks, simultaneous circulation of all four DENV serotypes, and the occurrence of epidemic cycles every 2-3 years. In Colombia, the largest dengue epidemic was recorded in 2010, with >150,000 confirmed cases and 217 deaths [3]. Moreover, during 2007–2017, 609,228 cases of dengue were reported, of which 119,888 (19.7%) occurred in the Caribbean Region, specifically in the departments of Atlantic, Cesar, Córdoba, Sucre, Bolívar, Guajira, Magdalena, and San Andrés y Providencia [4].

In addition to the occurrence of dengue, chikungunya and Zika viruses were recently introduced into Colombia. Regarding the chikungunya virus, the first locally-transmitted cases in Colombia were recorded in 2014 among the inhabitants of San Joaquin, municipality of Mahates, Department of Bolivar in the Caribbean region. Until 2017, 488,402 cases of chikungunya had been reported, of which 118,496 (24.3%) were reported in the departments of the Caribbean region [5]. Regarding the Zika virus, the first local outbreak of this disease occurred in 2015 in the municipality of Turbaco, Department of Bolivar. Up until 2017, 62,394 cases had been reported nation-wide, of which 6,288 (10.1%) were reported in the departments of the Caribbean Region [6].

The transmission of DENV, CHIKV, and ZIKV depends on three components—the host (in this case, humans), the virus, and the *A*. *aegypti* vector. Activities related to the prevention and control of these arboviruses in Colombia have predominantly focused on *A. aegypti* via community-directed educational campaigns for the elimination of mosquito breeding sites, the application of biological insecticides to larval habitats (in particular *Bacillus thuringiensis* var. *israellensis*), the use of insect growth regulators to treat larval habitats, and spraying of pyrethroid and organophosphate insecticides to control adult mosquitoes [7, 8]. Constant selection pressure by pyrethroid and organophosphate insecticides has resulted in the emergence of resistant *A. aegypti* populations in multiple areas of Colombia [9–15].

Resistance to insecticides in mosquitoes can be caused by the following mechanisms: behavioral modifications resulting in lessened likelihood of exposure, decreased penetration of the insecticide across the mosquito cuticle, alterations occurring at the insecticide target site within the mosquito, and increased detoxification (also referred to as metabolic resistance); the latter two mechanisms are the most frequently studied [16]. Target site alterations are most commonly caused by *kdr* mutations on the voltage-dependent sodium channel gene *para*, which is the target site for pyrethroids and DDT, or by mutations on the *Ace-1* gene (coding for the enzyme acetylcholinesterase), which is the target site for organophosphate and carbamate insecticides [17]. Metabolic resistance arises due to the increased activity or expression of genes coding for the main detoxifying enzymes including glutathione S-transferases, mixed-function oxidases, and esterases [16].

In Colombia, the insecticide susceptibility status of *A. aegypti* populations has been monitored for more than a decade. Since 2004, the National Insecticide Resistance Surveillance Network, headed by Colombia’s National Institute of Health, has evaluated approximately 170 populations of *A. aegypti* in 26 of the 32 departments in Colombia. The findings demonstrate variability in susceptibility to the insecticides temephos, λ-cyhalothrin, deltamethrin, permethrin, cyfluthrin, etofenprox, malathion, fenitrothion, pirimiphos-methyl, bendiocarb, and propoxur [12-14, 18-29]. Moreover, increased activity levels of insecticide-degrading enzymes, such as nonspecific esterases, mixed-function oxidases (MFOs), glutathione S-transferases (GSTs), and insensitive acetylcholinesterase (iAChE), have been observed in resistant populations [9–13, 26]. In addition, the *kdr* mutations V1016I [13, 30], F1534C [31], and V410L [15] associated with pyrethroid resistance have recently been detected.

Specifically in the Caribbean region, Maestre *et al.* [13] found variations in susceptibility to the organophosphates temephos, malathion, fenitrothion, and pirimiphos-methyl across *A. aegypti* populations. In addition, in the majority of the evaluated populations, resistance to the pyrethroids λ-cyhalothrin, deltamethrin, permethrin, and cyfluthrin was observed, with the exception of the population from Ciénaga (Magdalena), which remained susceptible. This study also reported the V1016I *kdr* mutation for the first time in Colombia.

Atencia *et al.* (2016) [31] found resistance to λ-cyhalothrin in populations of *A. aegypti* from the department of Sucre (Sincelejo) and reported the F1534C *kdr* mutation for the first time. Granada et al. (2018) [15] detected the V1016I and F1534C mutations in an *A. aegypti* populations from Riohacha (Guajira), with frequencies of 0.25 for V1016I and 0.71 for F1534C. Moreover, they reported the V410L mutation for the first time in Colombia, with allelic frequency of 0.30, in the populations from Riohacha. Notably, the *A. aegypti* mosquitoes in this population were resistant to λ-cyhalothrin.

The present study builds upon earlier work by further investigating the intensity and spatial extent of pyrethroid resistance in *A. aegypti* along the Caribbean coast of Colombia and links the frequency of *kdr* alleles and tri-locus *kdr* haplotypes to insecticide resistant phenotypes. To further understand the mechanisms of resistance, we also analyzed the activity levels of key detoxification enzyme groups.

## Materials and methods

### *A. aegypti* collections

*A. aegypti* were collected in the municipalities of Barranquilla (N 10° 57’ 10.622’’, W 75° 49’ 12.024’’) and Juan de Acosta (N 10° 49’ 44.731’’, W 75° 2’ 9.088’’) in the department of Atlantico; Cartagena (N 10° 24’ 55.416’’, W 75° 27’ 38.485’’) in the department of Bolivar; Valledupar (N 9° 56’ 55.068’’, W 73° 38’ 4.164’’) and Chiriguana (N 9° 21’ 41.27’’, W 73° 35’ 58.919’’) in the department of Cesar; and Monteria (N 8° 44’ 30.866’’, W 75° 52’ 0.433’’) in the department of Cordoba (Fig 1).

**Fig 1.**
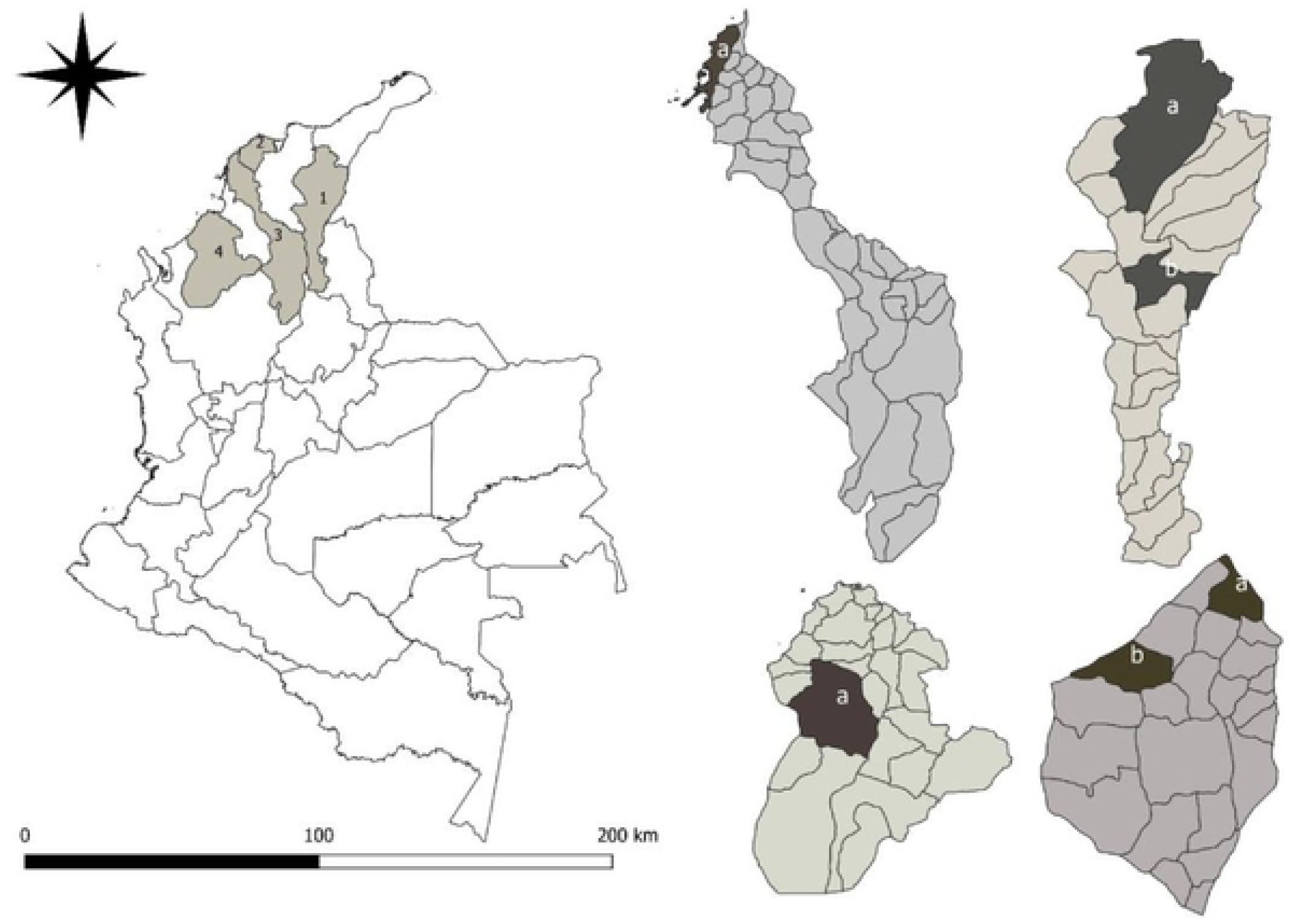
Collection sites of *Aedes aegypti* located in the Colombian Caribbean Region 1. Cesar: a) Valledupar, b) Chiriguana; 2. Atlantico: a) Barranquilla, b) Juan de Acosta; 3. Bolivar: a) Cartagena; 4) Cordoba: a) Monteria.

Immature stages were collected from habitats including tanks, pools, plastic/metallic cans, tires, animal water dishes, and flower vases located around houses. The specimens were reared to adults and maintained under controlled conditions of temperature (28°C ± 2°C), relative humidity (60% ± 10%), and photoperiod (12 h light:12 h dark) in the Public Health Laboratory of the department of Atlántico.

Upon emergence, male mosquitoes were fed with 10% sugar solution, and the females were fed with mouse (*Mus musculus)* blood every third day to obtain eggs of the F1 generation. Eggs were stored in an airtight plastic container, until they were hatched to obtain the adult mosquitoes used in the bioassays.

### Bioassays

Insecticide bioassays were performed following the methodologies described by the CDC [32] and WHO [33]. The pyrethroid insecticides and their concentrations were as follows: λ-cyhalothrin [10 µg/bottle (CDC) and 0.03% treated papers (WHO)], deltamethrin [10 µg/bottle (CDC) and 0.03% treated papers (WHO)], and permethrin [15 µg/bottle (CDC) and 0.25% treated papers (WHO)]. The technical grade insecticides (Chem Service®) used for the CDC bioassays were provided by the National Insecticide Resistance Surveillance Network of the Colombian National Institute of Health. The insecticide-impregnated papers used for the WHO bioassays were provided by Universiti Sains Malaysia.

For each population, 20-25 F1 generation, 3- to 5-day-old, unfed female *A. aegypti* were used in each bioassay replicate; as a control, the susceptible Rockefeller laboratory *A. aegypti* strain was used. Each bioassay consisted of four replications per insecticide for each population. The diagnostic time post-exposure was 30 min for the CDC bioassays and 24h for the WHO bioassays. Upon the completion of the diagnostic time, the living and dead specimens were classified as phenotypically resistant (R) or susceptible (S), and individually stored in 0.5-mL tubes with a hole in the lid and desiccated in tightly sealed bags containing silica gel. The bags containing the tubes were stored at −80°C for the subsequent detection of the V1016I, F1534C, and V410L *kdr* alleles.

In populations where resistance was detected via the CDC bioassay, resistance intensity was determined by conducting additional bioassays employing 2X the original insecticide concentration [33].

### Biochemical assays

Biochemical assays were conducted on F1 generation adults. One day post-emergence, 40 unfed female *A. aegypti* from each population were preserved at −80°C until processing. Individuals from the susceptible Rockefeller strain were used as controls. Mosquitoes were homogenized individually in 30 μl of distilled water for 5-10 seconds with an electric macerator and an additional 270 μl of distilled water was added for a final volume of 300 μl. Subsequently, each sample were centrifuged at 12,000 rpm for 60 seconds and aliquoted 10 μl for α, β, pNPA-esterases, 15 μl for GST, 20 μl for MFO and 25 μl for iAChE in triplicate in 96 well microplates. For the tests of mixed-function oxidases and acetylcholinesterase, the samples were transferred without being centrifuged. Enzyme activity levels were determined according to the methodology described by Valle *et al.* [34], which measures the optical densities at predetermined wavelengths to estimate the activity levels of MFO, iAChE, esterases, and GSTs. Total protein concentration was also determined for each individual to correct for differences in body sizes [35]. Results were read using an ELISA plate reader (Multiskan^TM^- Thermo Fisher Scientific®).

### Detection of *kdr* alleles

Real-time PCR was used to identify the V1016I, F1534C, and V410L *kdr* mutations. To estimate the allele frequencies in natural populations, 40-50 *A. aegypti* parental (F0) mosquitoes from each population were analyzed. To estimate associations between genotype and phenotype, all phenotypically resistant (R) and 30 randomly selected susceptible (S) individuals were analyzed per insecticide per population.

DNA was extracted from individual mosquitoes using the Quanta Biosciences Extracta^TM^ Kit. Individual mosquitoes were placed in sterile 0.2-mL tubes and 25 µL extraction buffer was added to each tube, followed by an incubation at 95°C for 30 min in a C1000 Bio-Rad CFX 96 Touch^TM^ Real- Time System thermocycler. At the end of the incubation, 25 µL of stabilization buffer was added. DNA was quantified using a NanoDrop^TM^ 2000/2000c spectrophotometer (ThermoFisher Scientific). PCR reactions were performed in a Bio-Rad C1000 CFX96 Real-Time System thermocycler. Genotype was determined by analyzing the melting curves of the PCR products. The V1016I mutation was amplified following the methodology described by Saavedra-Rodríguez *et al.* [36], using a final reaction volume of 20 µL, containing 6 μL of ddH_2_O, 10 μL of iQ^TM^ SYBR® Green Supermix (Bio-Rad), 1 μL of each of the V1016f, I1016f, and I1016r primers, and 1 μL of DNA template (Table 1). The cycling conditions were as follows: an initial denaturation at 95°C for 3 min followed by 40 cycles of: 95°C for 10 s, 60°C for 10 s, and 72°C for 30 s; and a final extension at 95°C for 10 s. The melting curves were determined by a denaturation gradient from 65°C to 95°C with an increase of 0.2°C every 10 seconds.

**Table 1.**
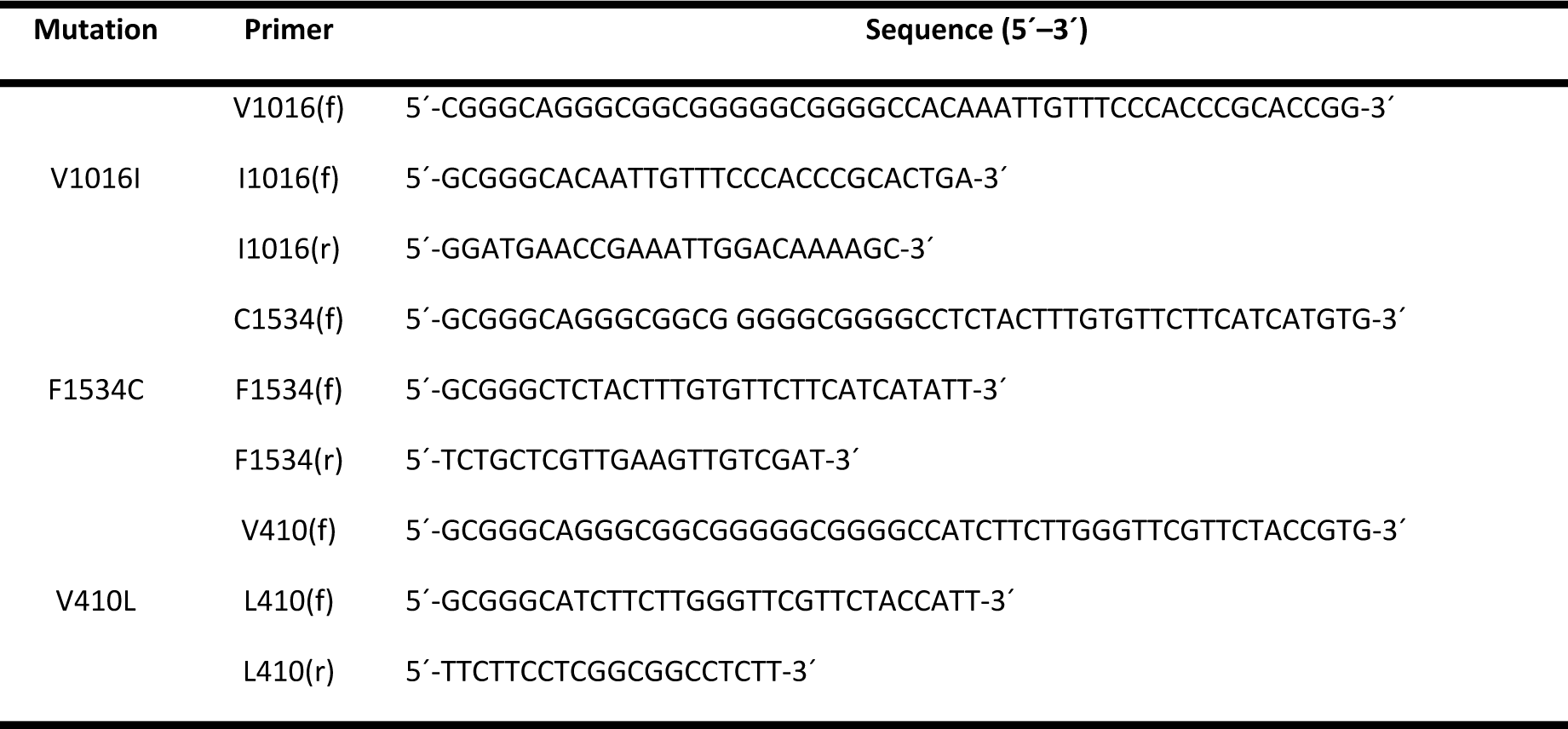
Primer sequences used for detecting *kdr* alleles

The F1534C mutation was detected following the methodology described by Yanola *et al.* [37], using a final reaction volume of 20 μL comprised of 7.15 μL of ddH_2_O, 9 μL of iQ^TM^ SYBR® Green Supermix (Bio-Rad), 0.6 μL of each of the F1534f and C1534r primers, 0.65 μL of the C1534f primers, and 2 μL of DNA template (Table 1). The cycling conditions were as follows: an initial denaturation at 95°C for 3 min followed by 37 cycles of: 95°C for 10 s, 57°C for 30 s, and 72°C for 30 s; and a final extension at 95°C for 10 s. The melting curves were determined by a denaturation gradient from 65°C to 95°C with an increase of 0.5°C every 5 s.

The V410L mutation was detected following the methodology described by Haddi *et al.* [38], using a final reaction volume of 21 μL comprised of 9.6 μL of ddH_2_O, 10 μL of iQ^TM^ SYBR® Green Supermix (Bio-Rad), 0.1 μL of each of the L410f and V410f primers, 0.2 μL of the L410r primer, and 1.0 μL of DNA template (Table 1). The cycling conditions were as follows: an initial denaturation at 95°C for 3 min followed by 39 cycles of: 95°C for 10 s, 60°C for 10 s, and 72°C for 30 s; and a final extension at 95°C for 10 s. The melting curves were determined by a denaturation gradient from 65°C to 95°C with an increase of 0.2°C every 10 s.

Each mosquito was analyzed in duplicate. For all assays for each mutation, three positive controls were included: a wild-type homozygote, a homozygote mutant, and a heterozygote. All assays also included a negative control consisting of master mix without DNA template.

### Data analysis

#### Bioassays

Mortality was scored at the diagnostic time per insecticide per population. Populations were categorized according to the WHO criteria [33], whereby 98%–100% mortality indicates susceptibility, 90%–97% suggests resistance is developing and <90% mortality indicates resistance.

#### Biochemical assays

Absorbance values were entered into Excel databases to calculate the average and standard deviation for each mosquito. To express the absorbance values in terms of enzymatic activity, data regarding the homogenate volume of each mosquito, total protein content for each mosquito, and units of activity for each enzyme group were calculated according to the protocol described by Valle *et al.* [34]. The cutoff value for the susceptible Rockefeller strain was determined based on the 99^th^ percentile of absorbance, and the percentage of individuals from the field strains with activity levels that exceeded this cutoff value were classified according to the criteria proposed by Montella *et al.* [39]: <15% unaltered, 15%–50% altered, and >50% highly altered.

After determining the activity levels for each enzyme group, an analysis of variance was performed, followed by Tukey’s multiple comparison test, with the significance level set at *p* ≤ 0.05, to identify populations with any statistically significant differences as compared to the Rockefeller reference strain.

#### Allelic and genotypic frequencies of the V1016I, F1534C, and V410L mutations

Results were obtained using Bio-Rad’s Precision Melt Analysis Software^TM^ and were interpreted as follows. For the V1016I mutation, a melting peak at 77°C corresponded to a mutant homozygote (I/I), a peak at 82°C corresponded to a wild-type homozygote (V/V), and peaks at both 77°C and 82°C corresponded to a heterozygote (V/I). For the F1534C mutation, a peak at 82°C corresponded to a mutant homozygote (C/C), a peak at 78°C corresponded to a wild-type homozygote (F/F), and peaks at both 78°C and 82°C corresponded to a heterozygote (F/C). For the V410L mutation, a peak at 80°C corresponded to a mutant homozygote (L/L), a peak at 83°C corresponded to a wild-type homozygote (V/V), and peaks at both 80°C and 83°C corresponded to a heterozygote (V/L).

From the parental mosquitoes (F0), the population-level allele frequencies for I1016, C1534, and L410 were calculated using Eq (1) as follows

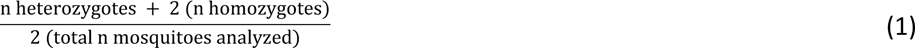

The genotypic frequencies for V_1016_/V_1016_, F_1534_/F_1534_, V_410_/V_410_, I_1016_/I_1016_, C_1534_/C_1534_, L_410_/L_410_, V_1016_/I_1016_, F_1534_/C_1534_, V_410_/L_410_ were calculated using Eq (2)

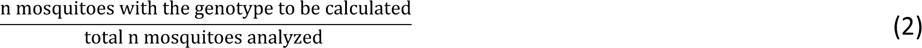

The Hardy–Weinberg principle was tested, as shown in Eq (3)

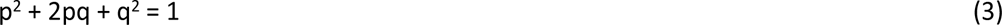

where p is the number of wild-type homozygotes, pq is the frequency of heterozygotes, and q is the frequency of mutant homozygotes.

Expected wild-type V_1016_/V_1016_, F_1534_/F_1534_, V_410_/V_410_ homozygotes = p^2^ (n) Expected V_1016_/I_1016_, F_1534_/C_1534_, V_410_/L_410_ heterozygotes = 2pq (n) Expected mutant I_1016_/I_1016_, C_1534_/C_1534_, L_410_/L_410_ homozygotes = q^2^ (n) The Chi square test was used to determine whether the populations were in Hardy–Weinberg equilibrium, as shown in Eq (4):

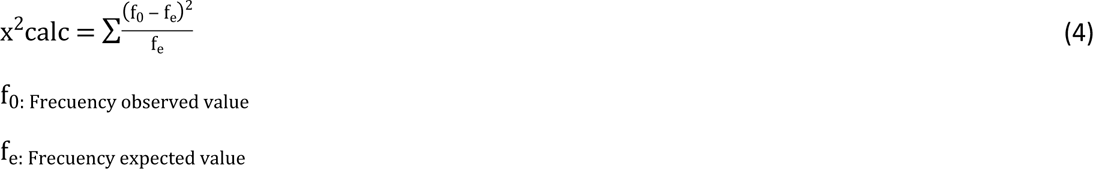

If the calculated value of χ^2^ was < tabulated χ^2^ (1 gl) = 3.84 and P < 0.05, the H_0_ that the study population was in Hardy–Weinberg equilibrium was accepted; otherwise, if the calculated χ^2^ was ≥ tabulated χ^2^, the H_a_ that the study population was not in Hardy–Weinberg equilibrium was accepted. In addition, the coefficient of endogamy was calculated using Eq (5) as follows:

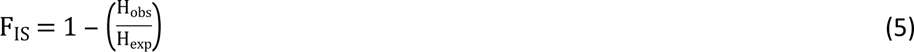

where, H_obs_ is the number of observed heterozygotes and H_exp_ is the number of expected heterozygotes; if F_IS_ was significantly higher than 0, an excess of homozygotes was considered, and if F_IS_ was significantly less than 0, an excess of heterozygotes was considered in the population, with a significance of P < 0.05. In addition, the frequencies of tri-locus genotypes were determined in the study populations.

#### Association of *kdr* mutations with pyrethroid resistance

The association between resistant and susceptible phenotypes and their *kdr* genotypes was tested using contingency tables, and the relationship between phenotype and tri-locus genotype was tested using the statistical software programs OpenEpi version 3.0 (https://www.openepi.com/TwobyTwo/TwobyTwo.htm) and GraphPad Prism version 8.1.

## Results

### Bioassays

A total of 1732 adult female *A. aegypti* were tested in WHO bioassays for susceptibility to λ- cyhalothrin (n=564), deltamethrin (n=586), and permethrin (n=582). Resistance to λ-cyhalothrin and permethrin was detected in all six evaluated populations. Resistance was most frequent in Monteria with 43.3% mortality to λ-cyhalothrin and 24.0% mortality to permethrin. Cartagena was the least resistant, with mortalities of 86.4% to λ-cyhalothrin and 77.6% to permethrin. Susceptibility to deltamethrin was observed in the populations from Juan de Acosta (98% mortality) and Barranquilla (100% mortality), and possible development of resistance was detected in Valledupar (96.8% mortality) and Monteria (93.2% mortality). The populations from Cartagena (87.9% mortality) and Chiriguana (86.0% mortality) were found to be resistant to deltamethrin (Fig 2).

**Fig 2.**
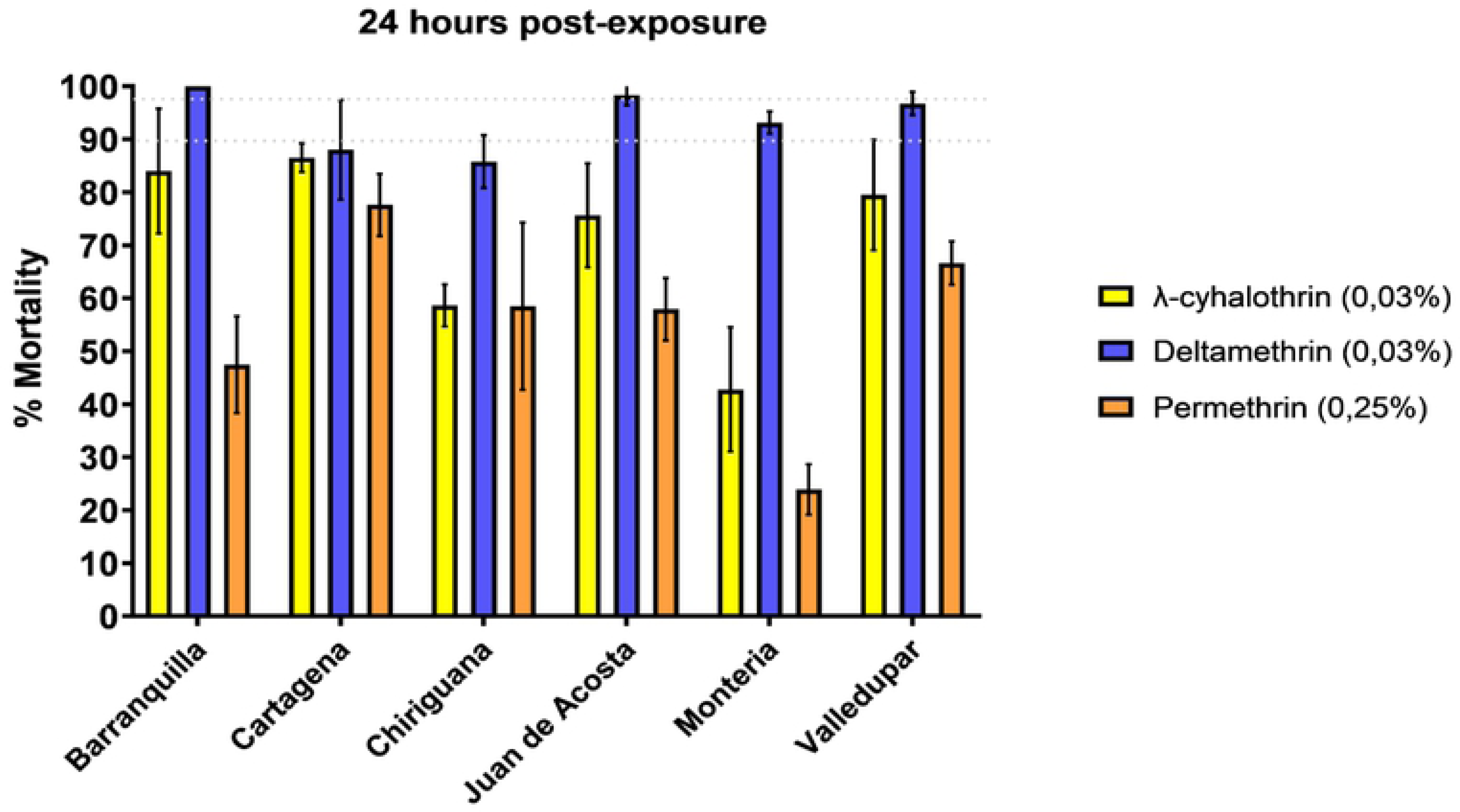
Mortality of the six populations of *A. aegypti* evaluated against diagnostic doses of pyrethroid insecticides following WHO bioassay methodology.

Additionally, a total of 1822 adult female *A. aegypti* were tested in CDC bioassays for susceptibility to λ-cyhalothrin (n=606), deltamethrin (n=608), and permethrin (n=608). Resistance to λ-cyhalothrin was detected in the populations from Barranquilla (79.6% mortality), Chiriguana (83.5% mortality), Juan de Acosta (71.6% mortality), and Monteria (35% mortality), whereas the populations from Cartagena (98.0% mortality) and Valledupar (100% mortality) were susceptible (Fig 3a). Susceptibility to deltamethrin was observed in all populations, with mortalities of 100% (Fig 3b). Resistance to permethrin was detected in the populations from Juan de Acosta (80.0% mortality), Monteria (69.0% mortality), and Barranquilla (64% mortality), and susceptibility was observed in the populations from Cartagena, Chiriguaná and Valledupar, with mortalities of 100% (Fig 3c). (Fig. 3). In populations where resistance intensity was assessed, 100% mortality at the diagnostic time was observed after exposure to twice the concentration (2X) of the recommended diagnostic dose for λ-cyhalothrin (20 μg/bottle) and permethrin (30 μg/bottle) (Table 2).

**Fig 3.**
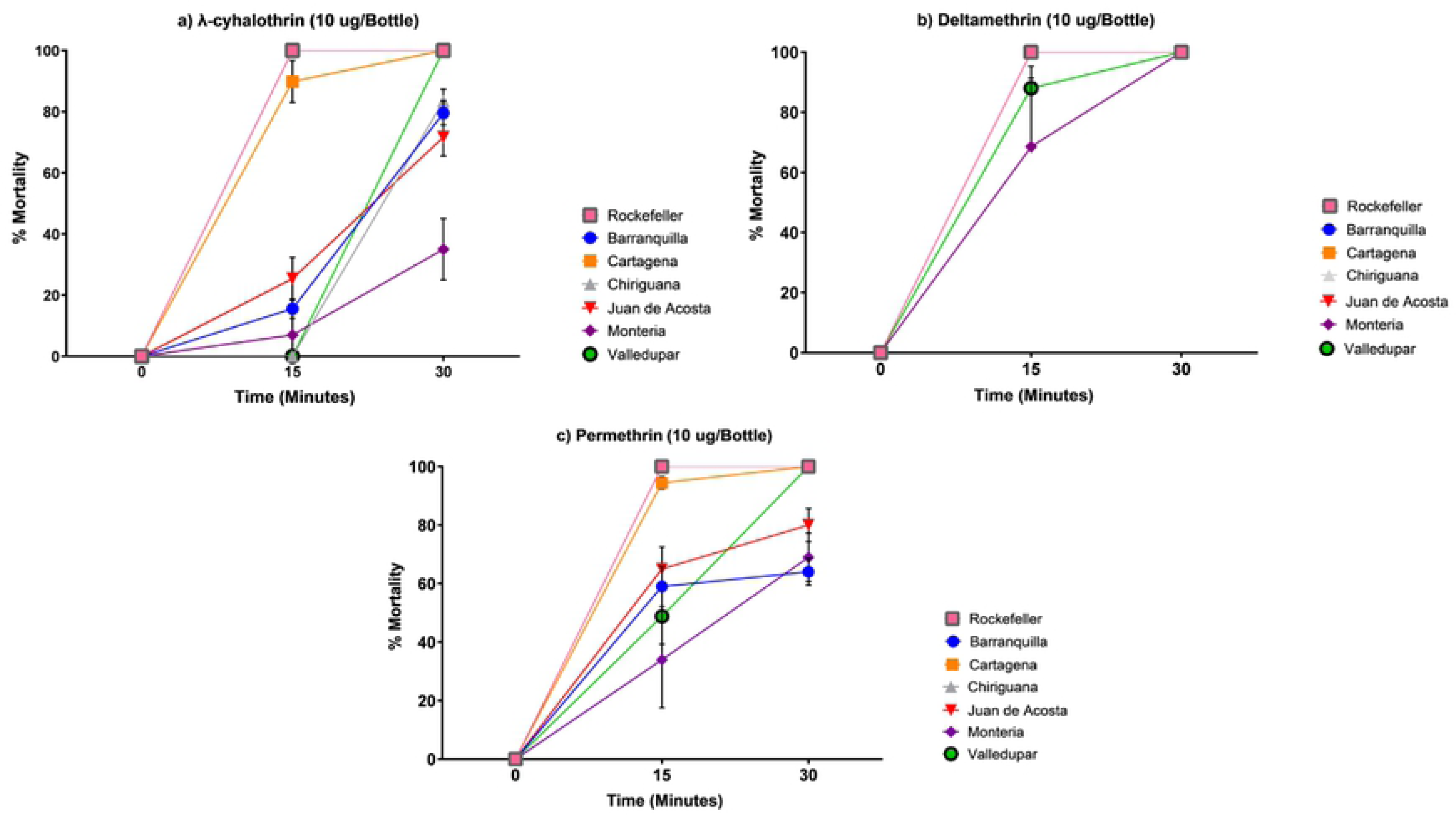
Mortality of the six populations of *A. aegypti* evaluated against diagnostic doses of pyrethroid insecticides following CDC bioassay methodology. a) λ-cyhalothrin (10ug/bottle), b) Deltamethrin (10ug/bottle, c) Permethrin (15ug/bottle).

**Table 2.** Mortality of *A. aegypti* exposed to 1X and 2X the diagnostic doses of λ-cyhalothrin and permethrin.

| Insecticide | Populations | 1× DD | | 2× DD | |
| --- | --- | --- | --- | --- | --- |
| | | 10 μg/bottle | | 20 μg/bottle | |
|  |  | n<sup>a</sup> | Mortality (%) | n | Mortality (%) |
| λ-cyhalothrin<br>(DD<sup>b</sup>: 10<br>μg/bottle, DT<sup>c</sup>: 30<br>min) | Barranquilla | 103 | 76.61 | 100 | 100 |
|  | Chiriguana | 103 | 83.49 | 100 | 100 |
|  | Juan de<br>Acosta | 102 | 71.56 | 100 | 100 |
|  | Monteria | 100 | 35.0 | 100 | 100 |

| Insecticide | Populations | 15 µg/bottle |  | 30 µg/bottle |  |
| --- | --- | --- | --- | --- | --- |
|  |  | N | Mortality (%) | n | Mortality (%) |
| **Permethrin** (DD: 15 µg/bottle, DT: 30 min) | Barranquilla | 100 | 64.0 | 100 | 100 |
|  | Juan de Acosta | 100 | 80.0 | 100 | 100 |
|  | Monteria | 100 | 69.0 | 100 | 100 |

a Total number of females evaluated, b Diagnostic dose, c Diagnostic time.

### Biochemical assays

Based on the classification criteria of Montella *et al.* [39], α-esterase enzyme levels were highly altered in the population from Monteria, where 79% of individuals exceeded the 99^th^ percentile of the Rockefeller reference population (Fig.4A). Similarly, β-esterase activity levels were highly altered in the population of Monteria (97%) and were also altered in the populations from Juan de Acosta (45%), Barranquilla (31%), Valledupar (27%) and Cartagena (12%) (Fig.4B), and pNPA-esterase activity levels were altered in the population from Juan de Acosta (14%) (Fig 4C). Highly altered MFO activity levels were detected in the populations from Juan de Acosta (92%), Monteria (97%), and Valledupar (88%) (Fig 4D). Altered GST activity levels were detected in the populations from Barranquilla (17%), Cartagena (24%), Juan de Acosta (44%), Monteria (34%) and Chiriguana (4%) (Fig. 4E). AChE activity remained unaltered in all populations evaluated (Fig. 4F). Overall, significant differences were observed between the mean activity levels of most enzyme groups between the field populations and the Rockefeller reference strain (p < 0.05) (Fig 4).

**Fig 4.**
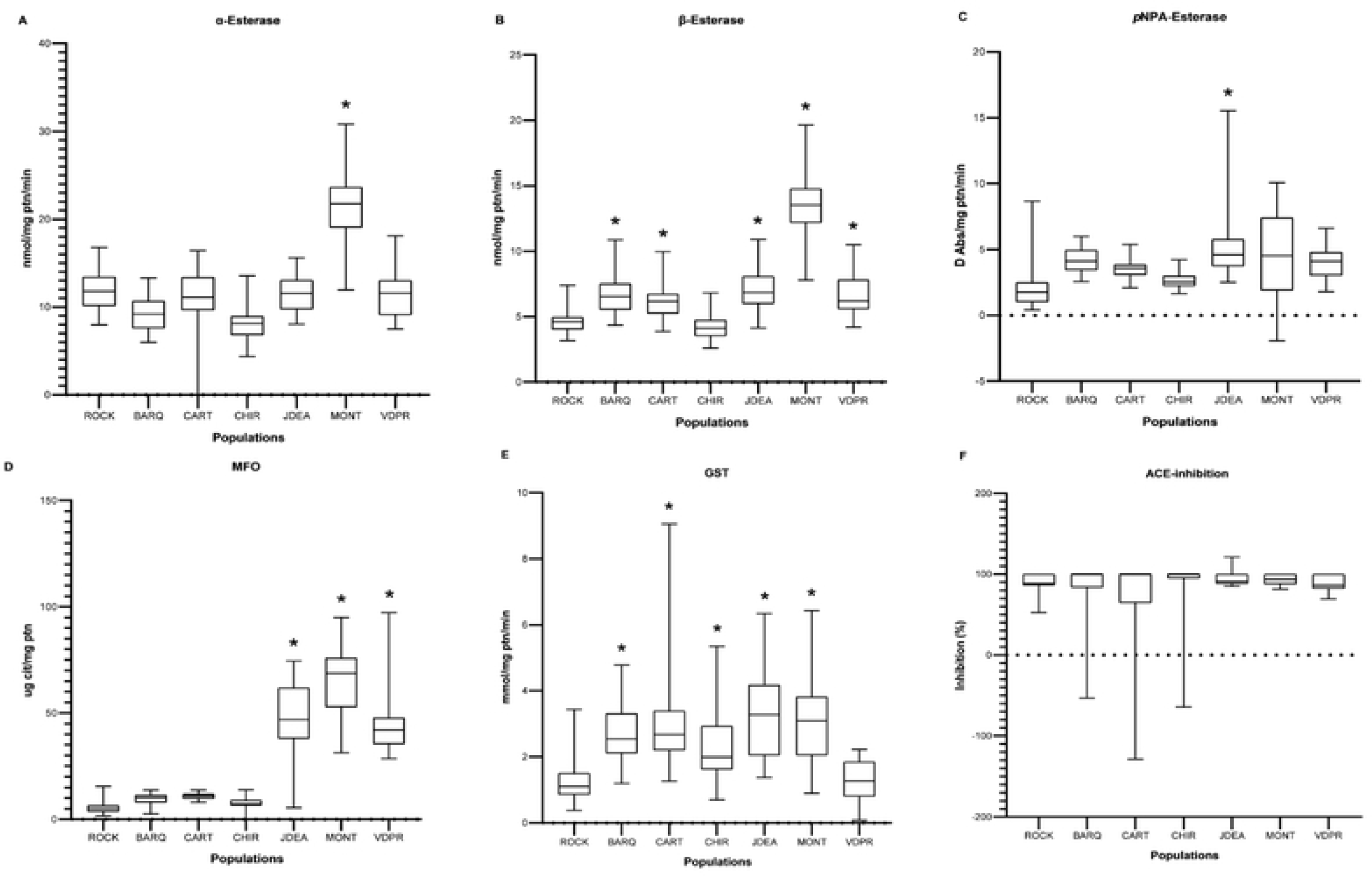
Box plots of enzymatic activity levels. *Aedes aegypti* populations with elevated enzymatic activity compared to the Rockefeller strain are marked with (*). (A). α-esterases, (B). β-esterases, (C). pNPA- esterases, (D). mixed-function oxidases (MFO), (E). glutathione-S-transferases (GSTs), and (F). insensitive acetylcholinesterase (iAChE). ROCK: Rockefeller; BARQ: Barranquilla-, CART: Cartagena; CHIR: Chiriguana; JDEA: Juan de Acosta; MONT: Monteria and VDPR: Valledupar.

### *kdr* allele frequencies

All three *kdr* mutations were detected in all the populations evaluated. Regarding the V1016I mutation, all three genotypes (VV_1016_, VI_1016_, and II_1016_) were detected in each field population. The mutant allele I1016 was the most prevalent in the population from Monteria, with a frequency of 0.70, and the least prevalent in the populations from Barranquilla and Valledupar, with a frequency of 0.15 in both. Regarding the F1534C mutation, all three genotypes (FF_1534_, FC_1534_, and CC_1534_) were detected in the populations from Barranquilla and Juan de Acosta, whereas only FC_1534_ and CC_1534_ were detected in Cartagena, Chiriguana, and Valledupar, with CC_1534_ predominating in all populations. It is noteworthy that the CC_1534_ genotype was fixed in the population from Monteria with a frequency of 1.0 (Table 3). In addition, the frequency of the C1534 mutant allele in the populations from Cartagena, Valledupar, and Chiriguana ranged between 0.94 and 0.97, but was 0.76 in the populations from Barranquilla and Juan de Acosta. Regarding the V410L mutation, all three genotypes (VV_410_, VL_410_, and LL_410_) were detected in each field population. The highest frequency of the L410 allele was detected in Montería with a frequency of 0.72, whereas the lowest was detected in Valledupar with a frequency of 0.05. For the other populations, the frequencies of the L410 allele ranged between 0.12 and 0.32 (Table 3).

**Table 3.**
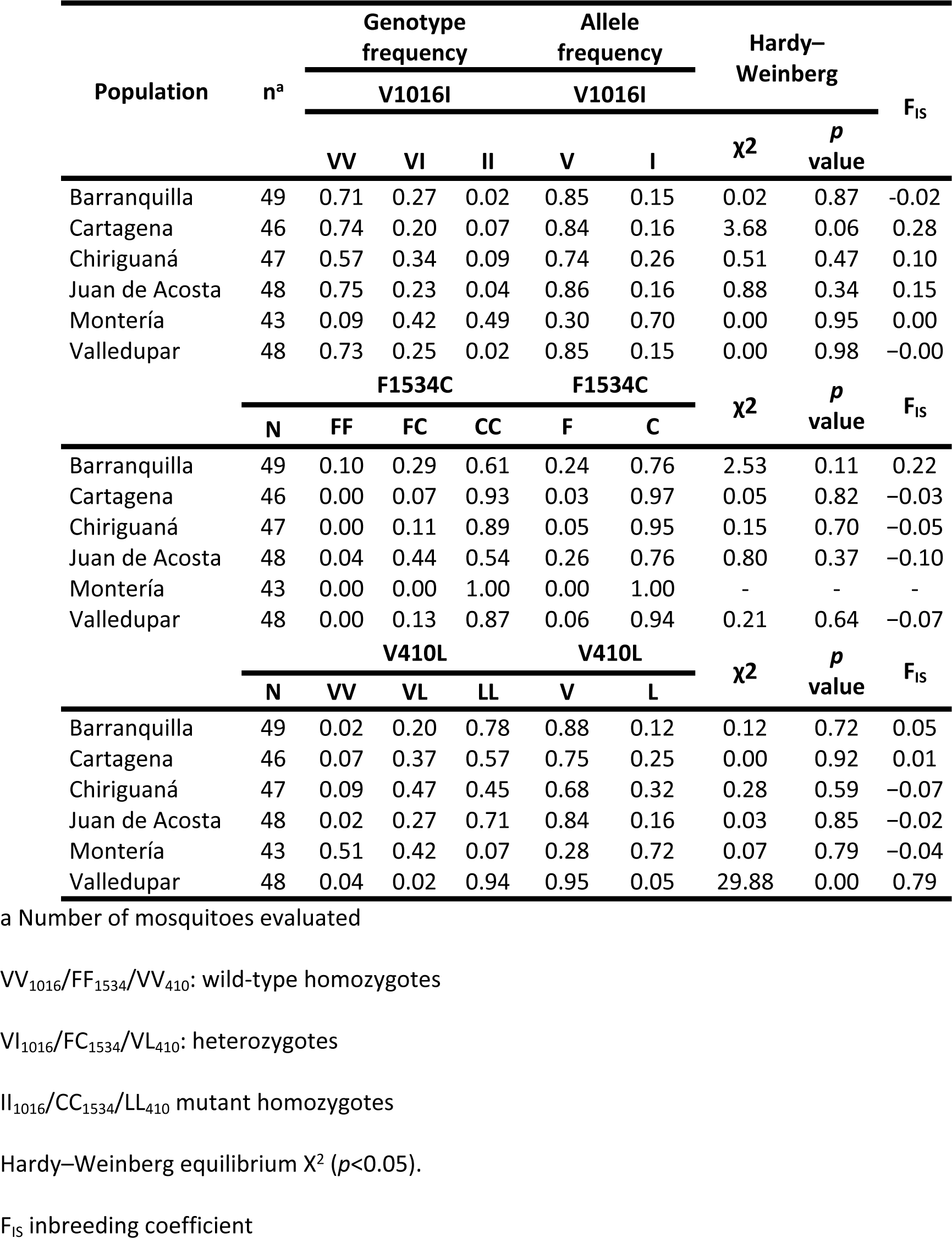
Genotype and allele frequencies of the V1016I, F1534C, and V410L *kdr* mutations in F0 *A. aegypti* females.

For loci 1016 and 1534, all genotypes were found to be in Hardy–Weinberg equilibrium. In the case of locus 410, the genotypes of most populations, except Valledupar, were in Hardy–Weinberg equilibrium (p < 0.05). When determining the inbreeding coefficients (F_IS_) for I1016, values < 0 were obtained for the populations from Barranquilla and Valledupar due to an excess of heterozygotes, in contrast to the populations from Cartagena, Chiriguana, Juan de Acosta, and Monteria, where values > 0 were recorded due to a deficiency of heterozygotes. For C1534, a generalized excess of heterozygotes was observed, with the exception of Barranquilla, where a deficiency of heterozygotes was observed. Similarly, for L410, the populations from Barranquilla, Cartagena, and Valledupar showed a deficiency of heterozygotes, in contrast to Chiriguana, Juan de Acosta, and Monteria, where an excess of heterozygotes was detected (Table 3).

Of the 27 combinations of tri-locus genotypes, 13 combinations were detected in 281 mosquitoes collected from the six evaluated populations. The triple homozygous wild-type genotype (VV_1016_, FF_1534_, and VV_410_) was detected only in the populations from Barranquilla and Juan de Acosta, with frequencies of 0.08 and 0.04, respectively, whereas the triple homozygous mutant genotype (II_1016_, CC_1534_, and LL_410_) was present in all populations except Valledupar, with frequencies between 0.02 (Barranquilla) and 0.49 (Monteria). Similarly, the triple heterozygous genotype (VI_1016_, FC_1534_, and VL_410_) was present only in Chiriguana and Juan de Acosta at low frequencies (0.02 and 0.06, respectively). The homozygous wild-type genotype for loci 1016 and 410 and homozygous resistant for locus 1534 (VV_1016_/CC_1534_/VV_410_) was most frequent in Barranquilla, Cartagena, Chiriguana, and Valledupar, with frequencies of 0.37, 0.54, 0.43, and 0.58, respectively; the exceptions were Juan de Acosta, where the most frequent genotype was homozygous wild-type for loci 1016 and 410 and heterozygous for locus 1534 (VV_1016_/FC_1534_/VV_410_) with a frequency of 0.33, and Monteria, where the most frequent genotype was the triple homozygous mutant (II_1016_/CC_1534_/LL_410_), with a frequency of 0.49 (Fig 5).

**Fig 5.**
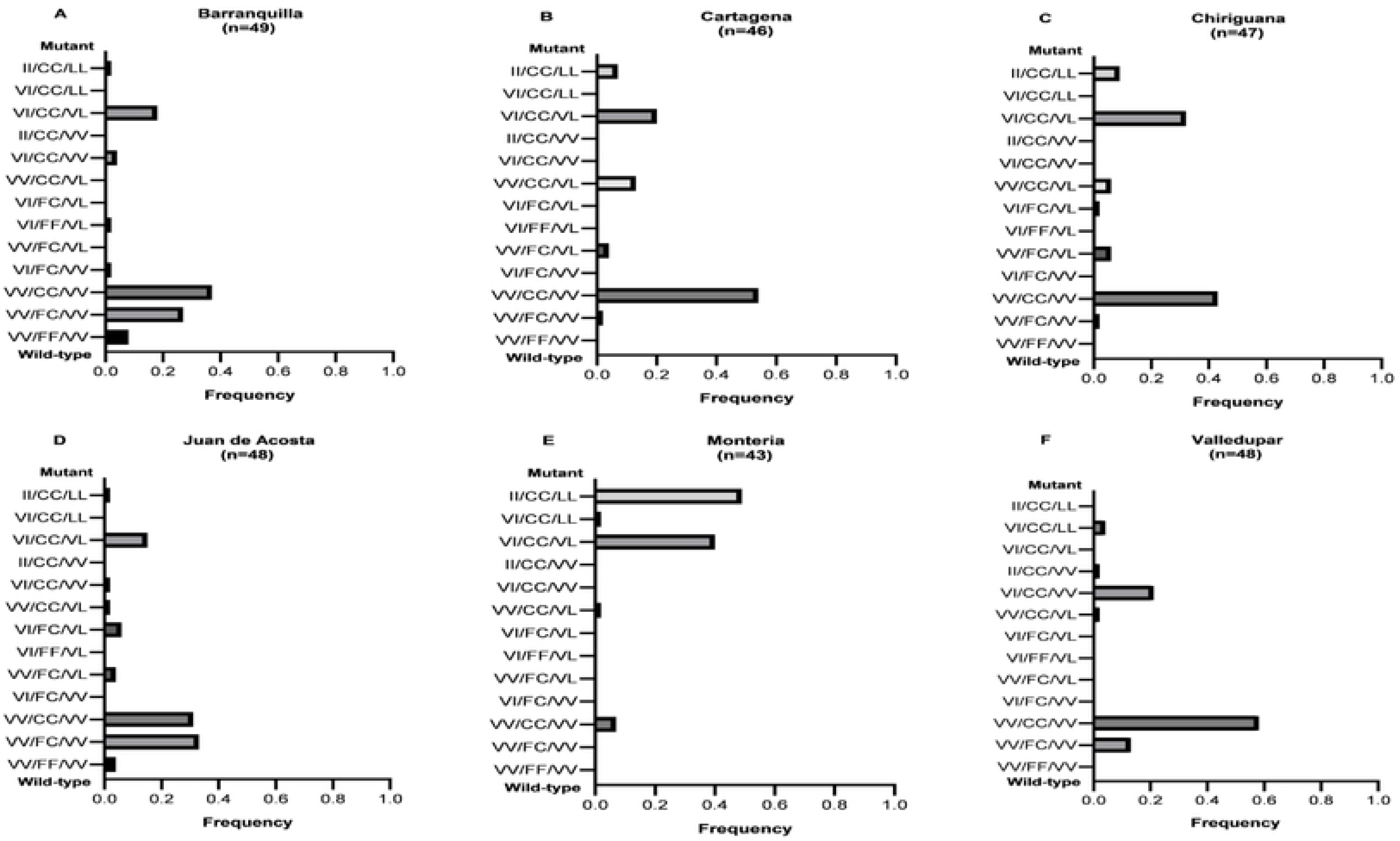
Frequencies of the 13 tri-locus genotypes present in F0 *A. aegypti* females. The order of the genotypes is 1016/1534/410. Mutant alleles: 1016 = I, 1534 = C, and 410 = L. The triple-mutant homozygous genotype is shown at the top and the triple-wild-type homozygous genotype at the bottom of each chart.

### Association of *kdr* alleles with phenotypic resistance to pyrethroids

Based on the results obtained with the mosquitoes exposed to insecticides in the WHO bioassays, a significant association (p < 0.05) was identified between the *kdr* alleles 1016I, 1534C, and 410L and resistance to λ-cyhalothrin in the populations from Juan de Acosta, Montería, and Valledupar. Similarly, an association was observed between the 1534C allele and resistance to deltamethrin in the populations of Chiriguana, Monteria, and Valledupar and between the 1016I and 410L alleles and resistance to deltamethrin in the population of Montería. A significant association (p < 0.05) was also detected between the 1016I, 1534C, and 410L alleles and resistance to permethrin in the populations from Chiriguana, Monteria, and Valledupar; between the 1534C allele and resistance to permethrin in Barranquilla, Cartagena, and Juan de Acosta; and between the 410L allele and permethrin resistance in Juan de Acosta (Tables 4–6).

**Table 4.**
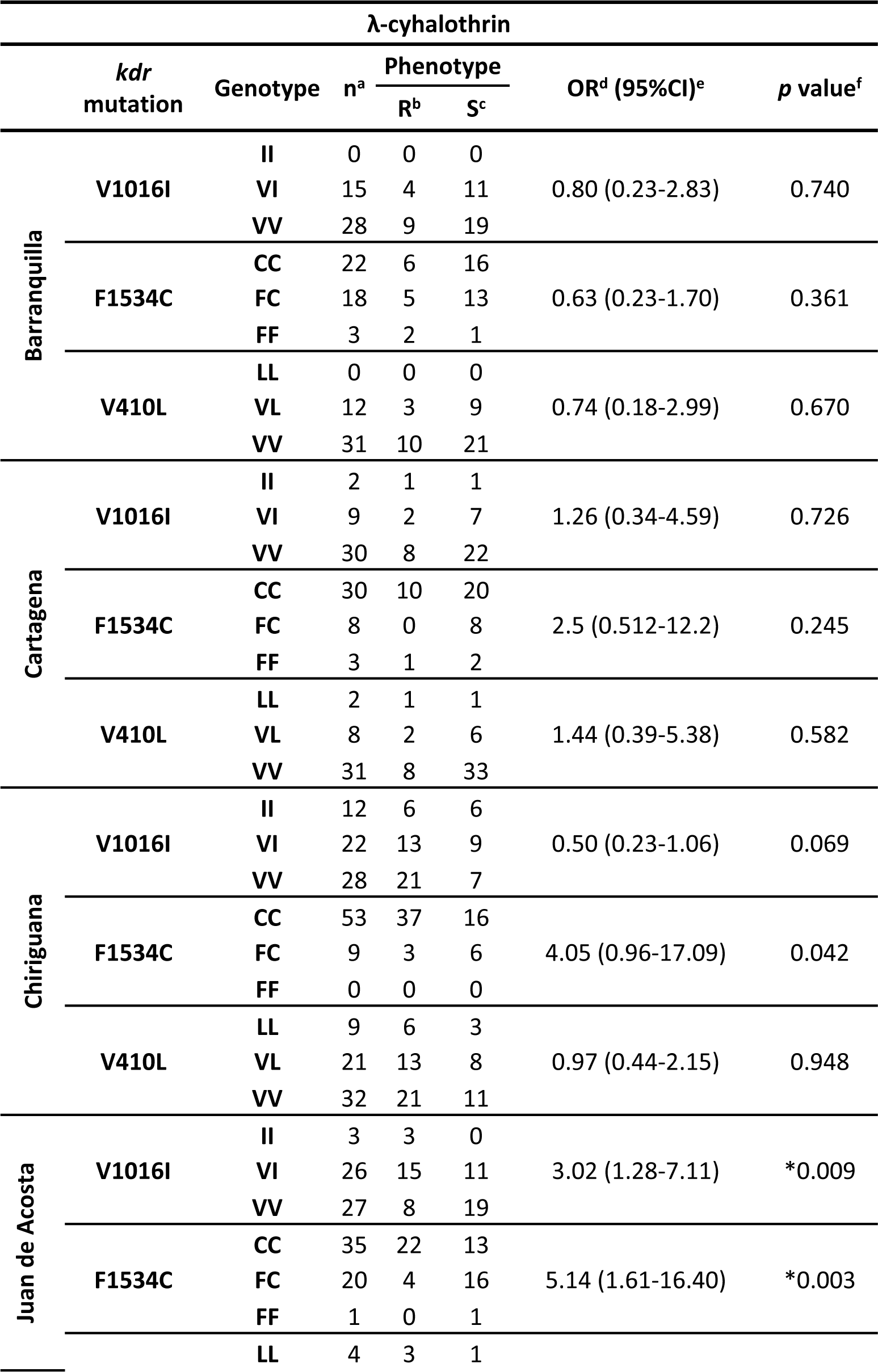
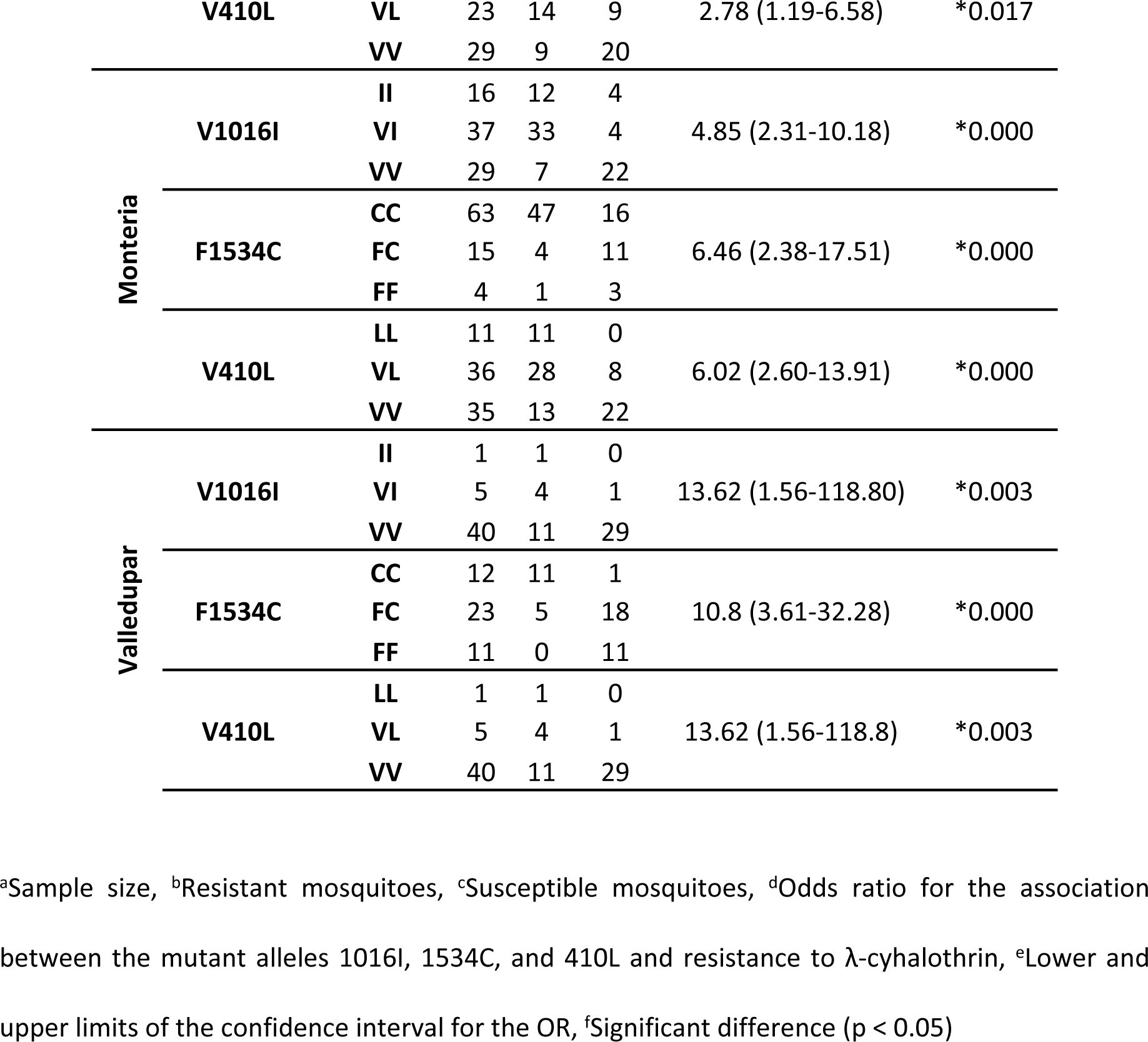
Association between 1016I, 1534C, and 410L alleles and resistance to λ-cyhalothrin in adult *A. aegypti* in WHO bioassays.

**Table 5.**
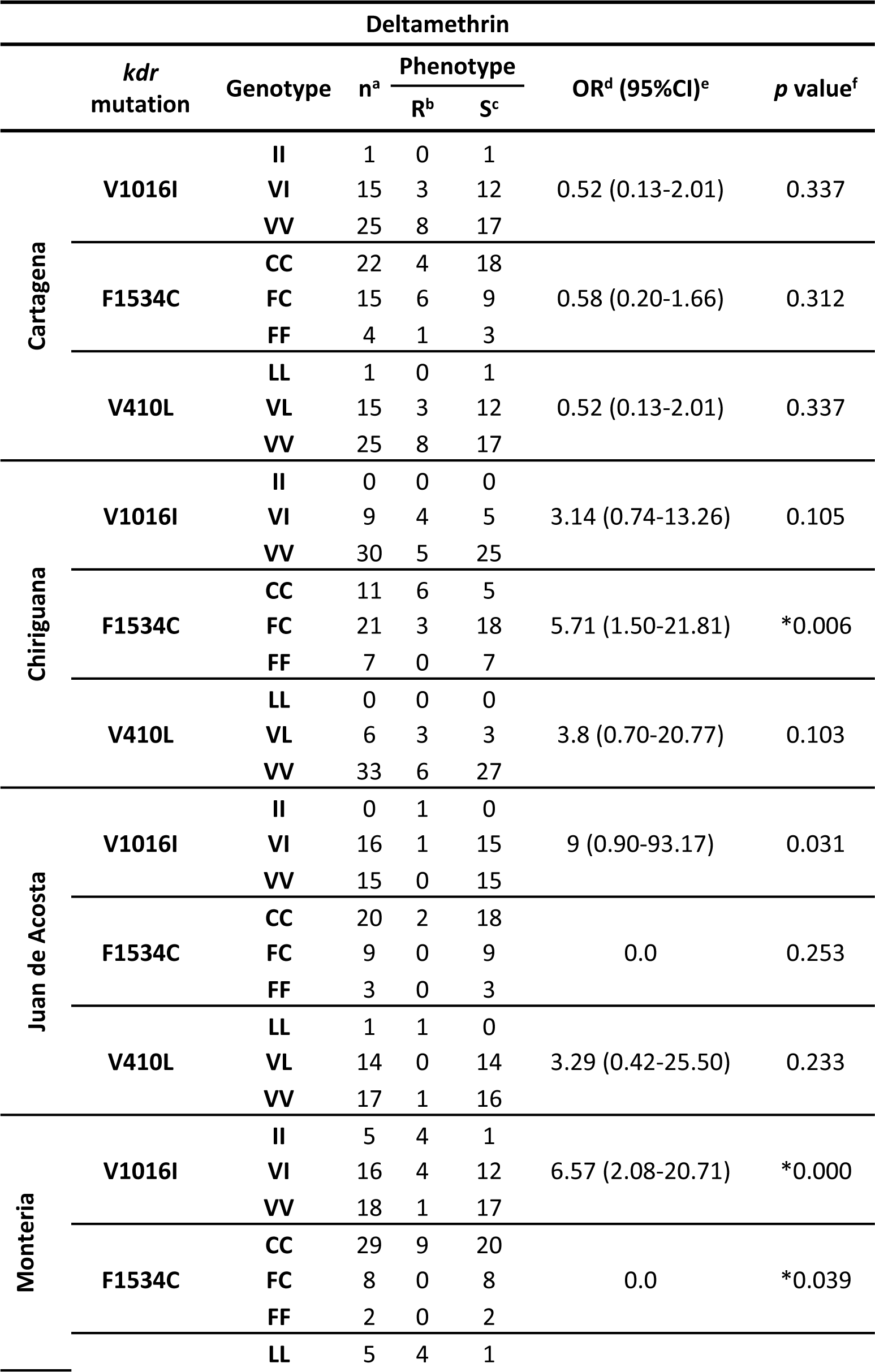
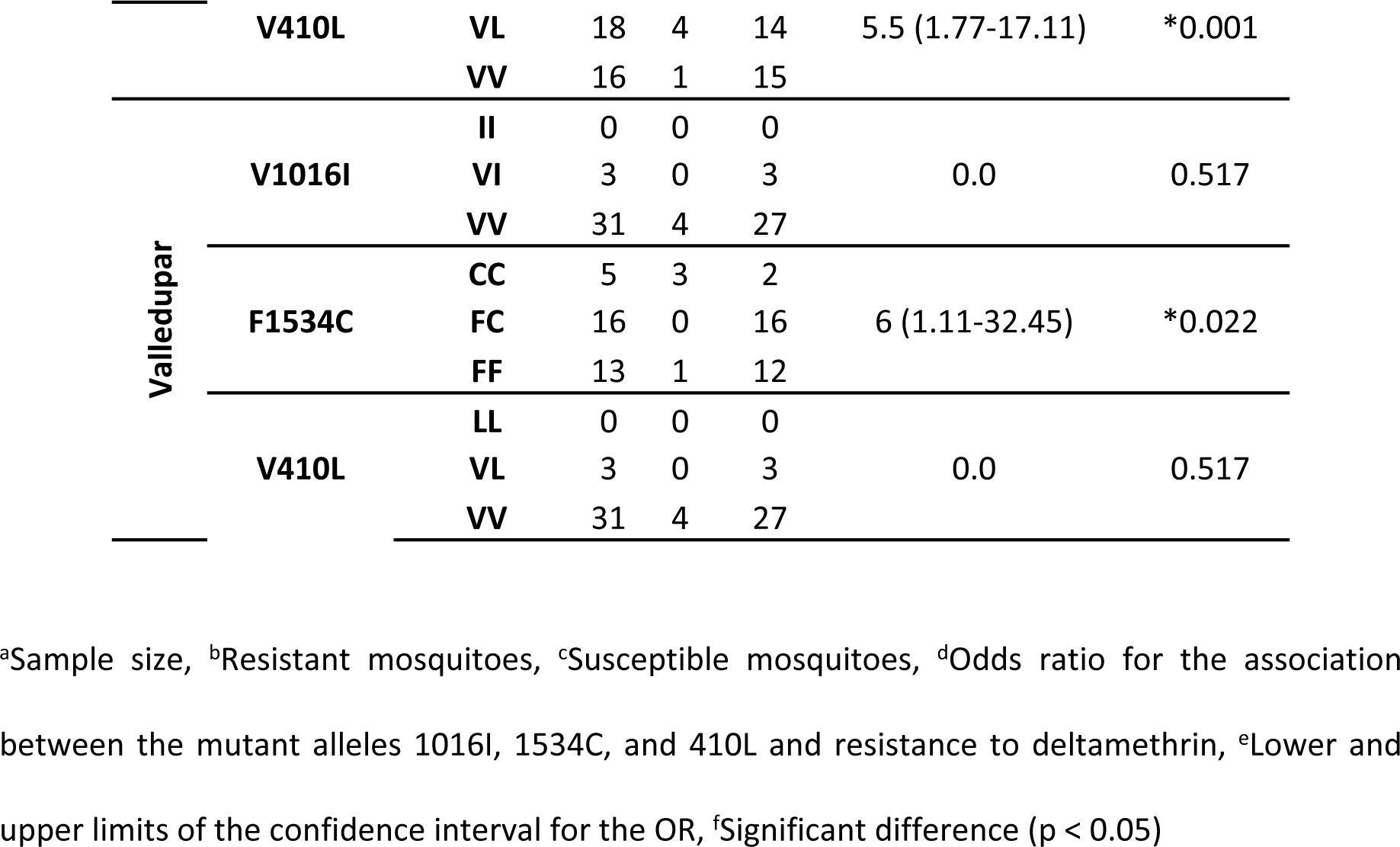
Association between 1016I, 1534C, and 410L alleles and resistance to deltamethrin in adult *A. aegypti* in WHO bioassays.

**Table 6.**
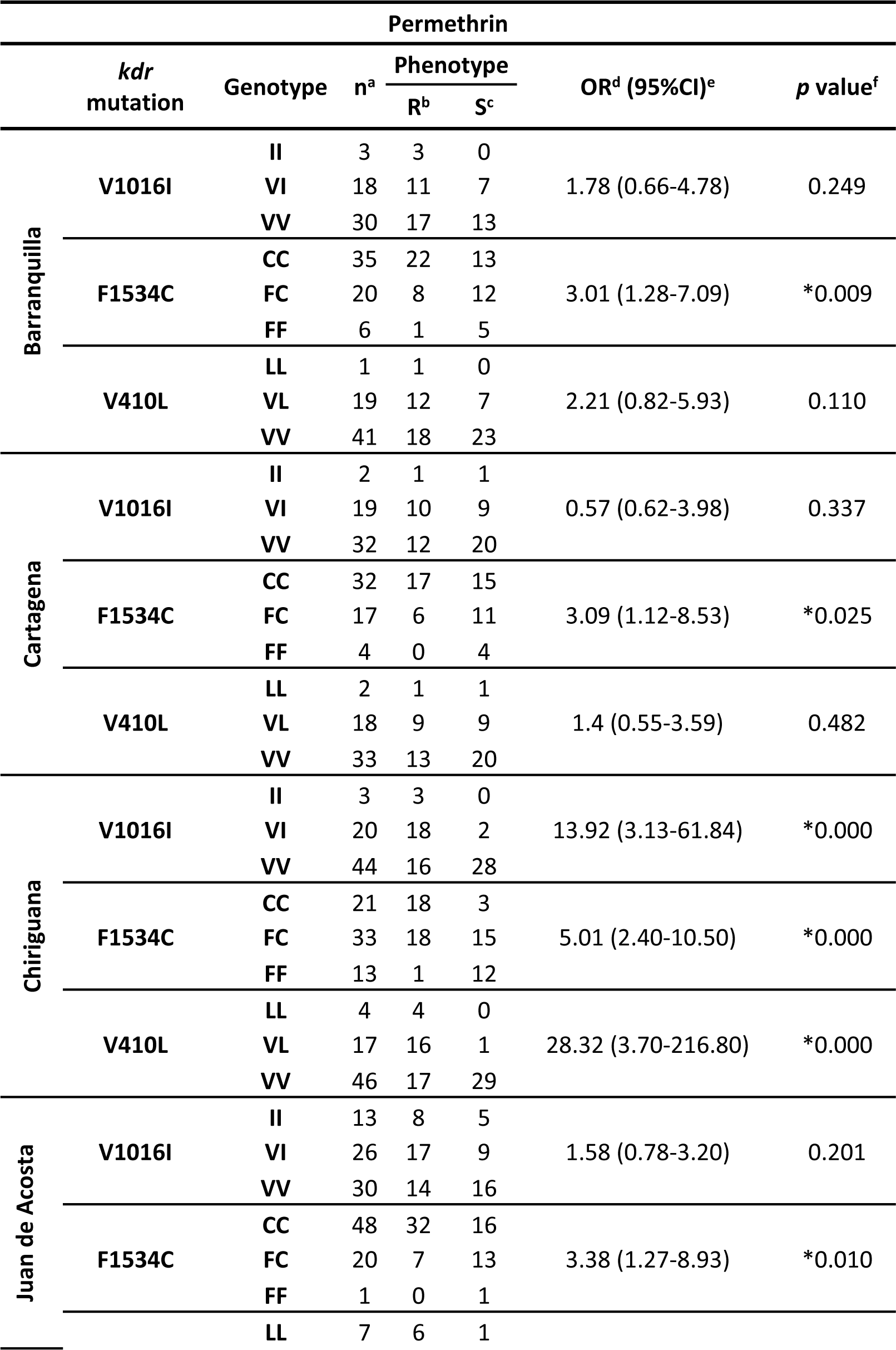
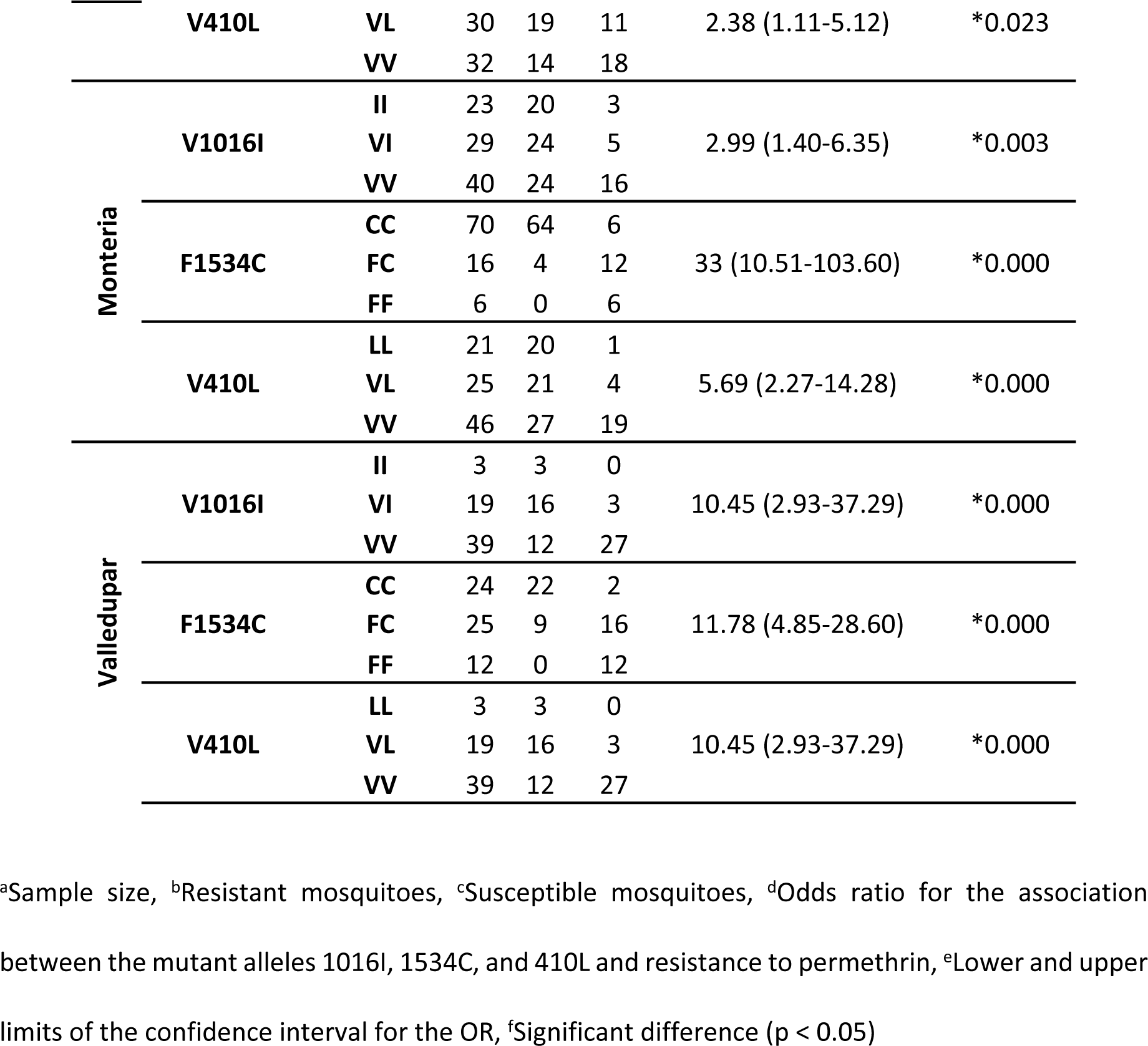
Association between 1016I, 1534C, and 410L alleles and resistance to permethrin in adult *A. aegypti* in WHO bioassays.

Less association was detected between *kdr* alleles and the observed phenotype in the CDC bioassays. A significant association (p < 0.05) between the 1534C allele and resistance to λ- cyhalothrin was detected in the population from Barranquilla and between the 1016I and 410L alleles and resistance to permethrin in the population from Montería. Despite the resistance to pyrethroids detected with the CDC bioassays in the populations from Chiriguana and Juan de Acosta, no significant associations were detected between *kdr* alleles and resistant phenotypes in these populations (Tables 7-8).

**Table 7.**
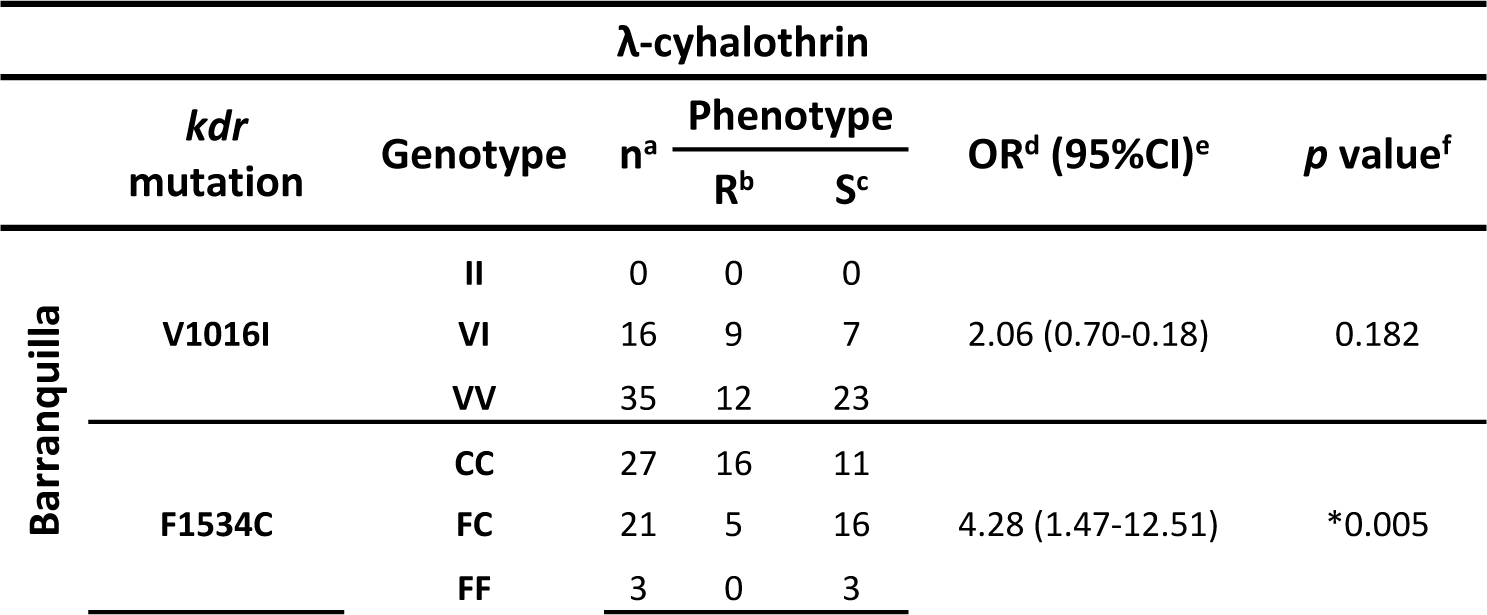
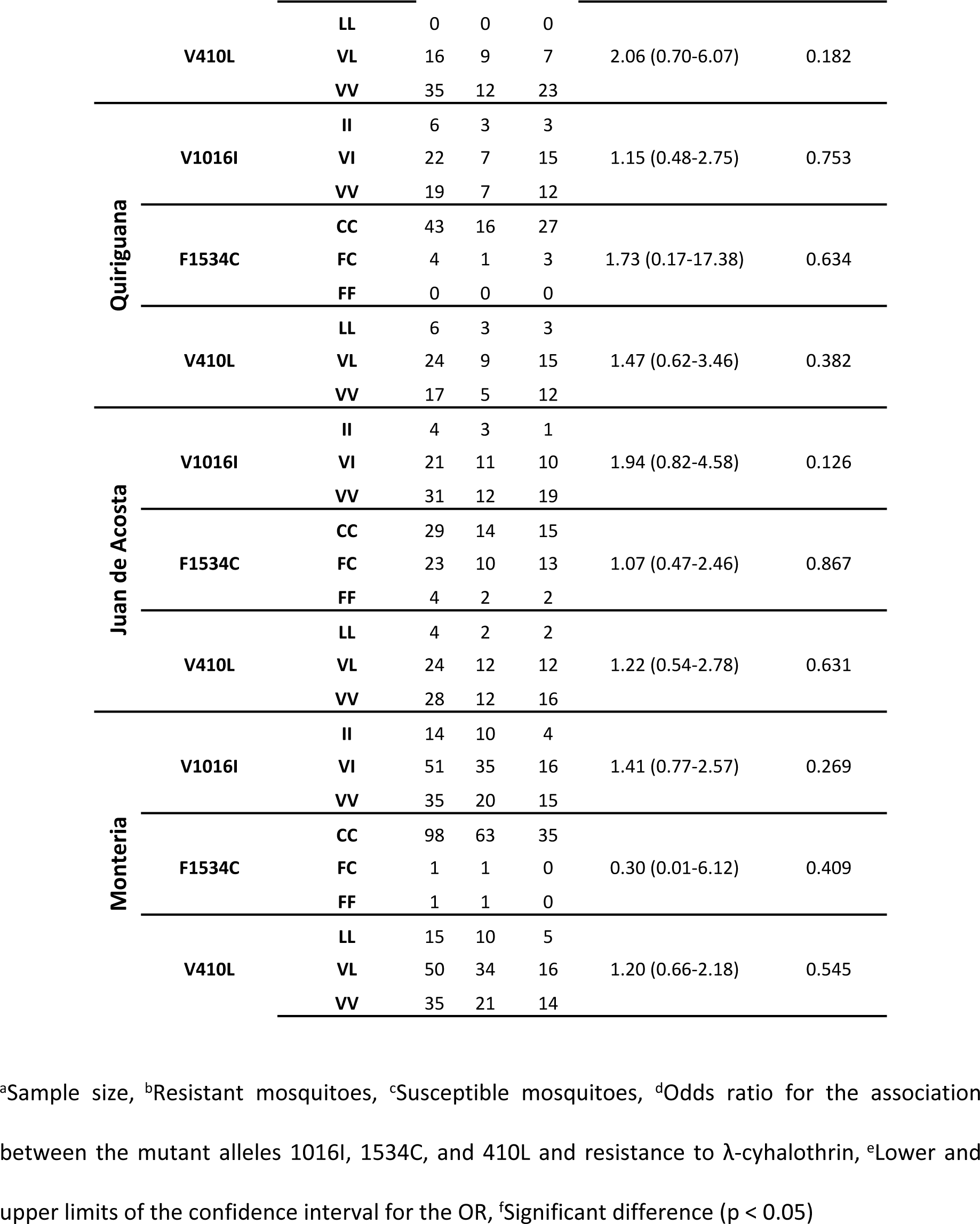
Association between 1016I, 1534C, and 410L alleles and resistance to λ-cyhalothrin in adult *A. aegypti* in CDC bioassays.

**Table 8.**
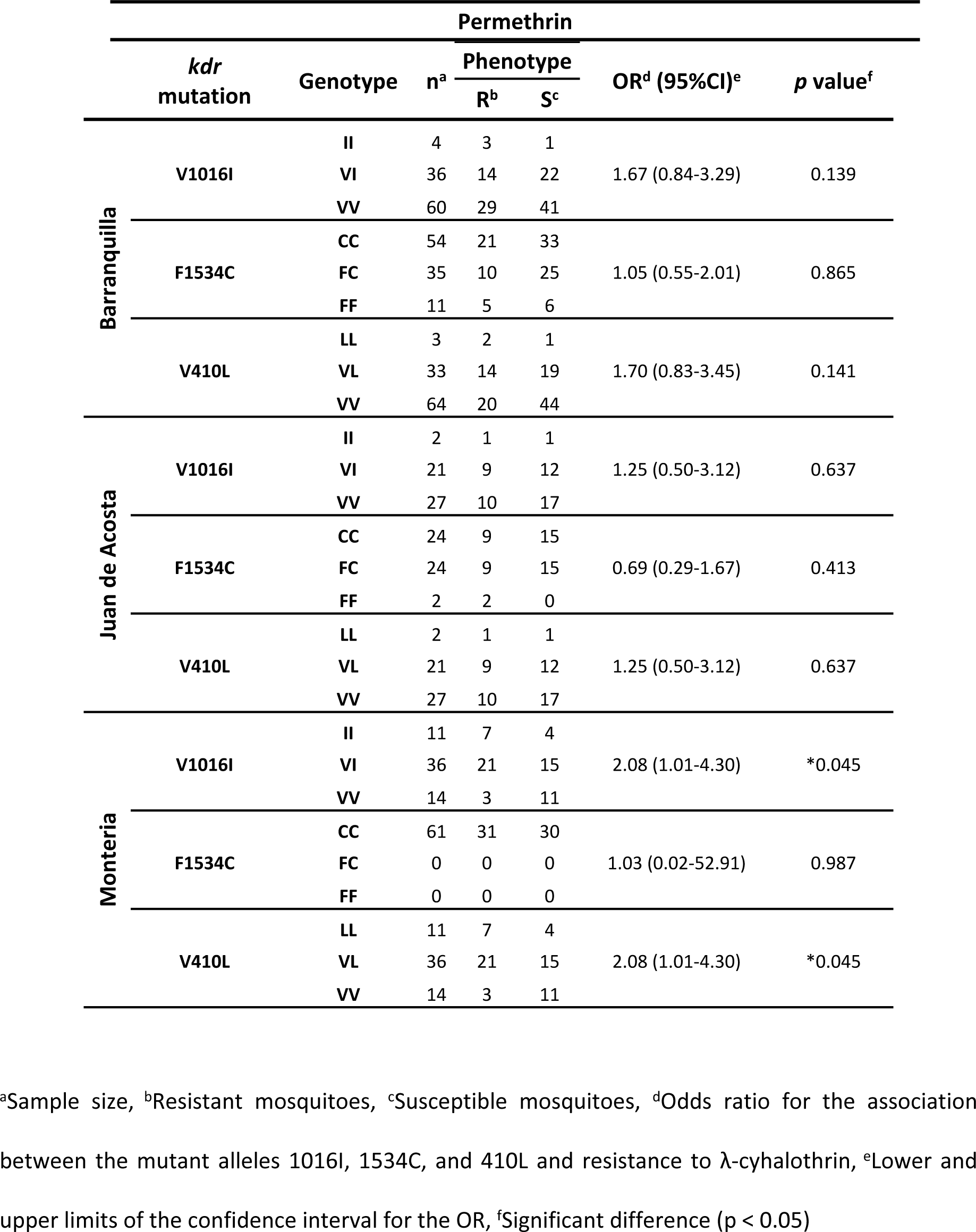
Association between 1016I, 1534C, and 410L alleles and resistance to permethrin in adult *A. aegypti* in CDC bioassays.

|  | *kdr*<br>mutation | Genotype | n<sup>a</sup> | Phenotype |  | OR<sup>d</sup> (95%CI)<sup>e</sup> | *p* value<sup>f</sup> |
| --- | --- | --- | --- | --- | --- | --- | --- |
|  |  |  |  | R<sup>b</sup> | S<sup>c</sup> |  |  |
| Barranquilla | V1016I | II | 4 | 3 | 1 | 1.67 (0.84-3.29) | 0.139 |
|  |  | VI | 36 | 14 | 22 |  |  |
|  |  | VV | 60 | 29 | 41 |  |  |
|  | F1534C | CC | 54 | 21 | 33 | 1.05 (0.55-2.01) | 0.865 |
|  |  | FC | 35 | 10 | 25 |  |  |
|  |  | FF | 11 | 5 | 6 |  |  |
|  | V410L | LL | 3 | 2 | 1 | 1.70 (0.83-3.45) | 0.141 |
|  |  | VL | 33 | 14 | 19 |  |  |
|  |  | VV | 64 | 20 | 44 |  |  |
| Juan de Acosta | V1016I | II | 2 | 1 | 1 | 1.25 (0.50-3.12) | 0.637 |
|  |  | VI | 21 | 9 | 12 |  |  |
|  |  | VV | 27 | 10 | 17 |  |  |
|  | F1534C | CC | 24 | 9 | 15 | 0.69 (0.29-1.67) | 0.413 |
|  |  | FC | 24 | 9 | 15 |  |  |
|  |  | FF | 2 | 2 | 0 |  |  |
|  | V410L | LL | 2 | 1 | 1 | 1.25 (0.50-3.12) | 0.637 |
|  |  | VL | 21 | 9 | 12 |  |  |
|  |  | VV | 27 | 10 | 17 |  |  |
| Monteria | V1016I | II | 11 | 7 | 4 | 2.08 (1.01-4.30) | *0.045 |
|  |  | VI | 36 | 21 | 15 |  |  |
|  |  | VV | 14 | 3 | 11 |  |  |
|  | F1534C | CC | 61 | 31 | 30 | 1.03 (0.02-52.91) | 0.987 |
|  |  | FC | 0 | 0 | 0 |  |  |
|  |  | FF | 0 | 0 | 0 |  |  |
|  | V410L | LL | 11 | 7 | 4 | 2.08 (1.01-4.30) | *0.045 |
|  |  | VL | 36 | 21 | 15 |  |  |
|  |  | VV | 14 | 3 | 11 |  |  |

<sup>a</sup>Sample size, <sup>b</sup>Resistant mosquitoes, <sup>c</sup>Susceptible mosquitoes, <sup>d</sup>Odds ratio for the association441 between the mutant alleles 1016I, 1534C, and 410L and resistance to λ-cyhalothrin, <sup>e</sup>Lower and442 upper limits of the confidence interval for the OR, <sup>f</sup>Significant difference ( $p < 0.05$ )

### Comparisons of tri-locus genotypes with resistance to pyrethroids

Of the 27 possible combinations of genotypes, 20 combinations of tri-locus genotypes were detected in the 918 mosquitoes phenotyped in WHO bioassays. The most common haplotypes were VV_1016_/CC_1534_/VV_410_ (n=233 mosquitoes, 25.4%), VV_1016_/FC_1534_/VV_410_ (n=198, 21.6%), and VI_1016_/CC_1534_/VL_410_ (n=187, 20.4%). Wild-type double homozygotes at loci 1016 and 410 in the presence of CC1534/FC1534 were significantly more likely to be phenotypically susceptible to deltamethrin (p < 0.05). Heterozygotes at both loci 1016 and 410 in the presence of CC1534 were significantly more likely to be resistant to λ-cyhalothrin and permethrin (p < 0.05) and susceptible to deltamethrin (p < 0.05) (Table 9).

**Table 9.** Tri-locus genotypes of phenotyped adult *A. aegypti* from the six study populations after WHO bioassay.

| tri-locus genotype |  |  |  |  |  |  |  |  |  |  |  |  |  |  |  |  |  |  |  |  |  |  |  |  |  |
| --- | --- | --- | --- | --- | --- | --- | --- | --- | --- | --- | --- | --- | --- | --- | --- | --- | --- | --- | --- | --- | --- | --- | --- | --- | --- |
| Insecticide | Phenotype | n<sup>c</sup> | II/CC/LL | II/FC/LL | VI/CC/LL | II/CC/VL | VI/FC/LL | VV/CC/LL | II/FC/VL | VI/CC/VL | II/CC/VV | VI/CC/VV | VV/CC/VL | II/FC/VV | VI/FC/VL | VV/FF/LL | VI/FF/VL | VV/FC/VL | II/FF/VV | VI/FC/VV | VV/CC/VV | VV/FF/VL | VI/FF/VV | VV/FC/VV | VV/FF/VV |
| λ cyhalothrin | R<sup>a</sup> | 158 | 22 | 0 | 0 | 1 | 0 | 0 | 0 | 56 | 0 | 7 | 0 | 0 | 7 | 0 | 0 | 0 | 0 | 0 | 47 | 0 | 1 | 14 | 3 |
|  | S<sup>b</sup> | 172 | 4 | 0 | 0 | 5 | 1 | 0 | 0 | 18 | 2 | 6 | 0 | 0 | 18 | 0 | 0 | 0 | 0 | 0 | 47 | 0 | 0 | 53 | 18 |
| Deltamethrin | R | 35 | 5 | 0 | 0 | 0 | 0 | 0 | 0 | 8 | 0 | 1 | 0 | 0 | 2 | 0 | 0 | 0 | 0 | 1 | 10 | 0 | 0 | 6 | 2 |
|  | S | 150 | 2 | 0 | 0 | 0 | 0 | 0 | 0 | 30 | 0 | 0 | 0 | 0 | 14 | 0 | 0 | 1 | 0 | 3 | 31 | 1 | 0 | 42 | 26 |
| Permethrin | R | 229 | 33 | 0 | 1 | 4 | 1 | 0 | 0 | 66 | 0 | 8 | 3 | 0 | 19 | 0 | 0 | 1 | 0 | 2 | 60 | 0 | 0 | 29 | 2 |
|  | S | 174 | 2 | 0 | 1 | 3 | 0 | 0 | 1 | 9 | 0 | 2 | 0 | 1 | 21 | 0 | 1 | 0 | 1 | 2 | 38 | 0 | 0 | 54 | 38 |
| Total |  | 918 | 68 | 0 | 2 | 13 | 2 | 0 | 1 | 187 | 2 | 24 | 3 | 1 | 81 | 0 | 1 | 2 | 1 | 8 | 233 | 1 | 1 | 198 | 89 |

<sup>a</sup>Resistant (living), <sup>b</sup>Susceptible (dead), <sup>c</sup>Total number of mosquitoes. The order of the genotypes is shown for loci 1016/1534/410. Resistant allele at locus 1016 = I, 1534 = C, 410 = L, triple-resistant
genotype II/CC/LL, triple-susceptible genotype VV/FF/VV. Significant differences between resistant and susceptible are shown in bold ( $p < 0.05$ ).

From the CDC bioassays, 15 combinations of tri-locus genotypes were observed in 465 mosquitoes assayed with λ-cyhalothrin and permethrin in Barranquilla, Juan de Acosta, and Monteria. Similar to the WHO bioassays, the most common haplotypes were VI_1016_/CC_1534_/VL_410_ (n=161, 34.6%) and VV_1016_/CC_1534_/VV_410_ (n=117, 25.2%). Wild-type double homozygotes at loci 1016 and 410 in the presence of CC1534/FC1534 were significantly more likely to be phenotypically susceptible to λ- cyhalothrin and permethrin (p < 0.05) (Table 10).

**Table 10.** Tri-locus genotypes of phenotyped adult *A. aegypti* after CDC bioassay.

| tri-locus genotype |  |  |  |  |  |  |  |  |  |  |  |  |  |  |  |  |  |  |  |  |  |  |  |  |  |
| --- | --- | --- | --- | --- | --- | --- | --- | --- | --- | --- | --- | --- | --- | --- | --- | --- | --- | --- | --- | --- | --- | --- | --- | --- | --- |
| Insecticide | Phenotype | n<sup>c</sup> | II/CC/LL | II/FC/LL | VI/CC/LL | II/CC/VL | VI/FC/LL | VV/CC/LL | II/FC/VL | VI/CC/VL | II/CC/VV | VI/CC/VV | VV/CC/VL | II/FC/VV | VI/FC/VL | VV/FF/LL | VI/FF/VL | VV/FC/VL | II/FF/VV | VI/FC/VV | VV/CC/VV | VV/FF/VL | VI/FF/VV | VV/FC/VV | VV/FF/VV |
| λ cyhalothrin | R<sup>a</sup> | 129 | 14 | 0 | 1 | 2 | 0 | 0 | 0 | 52 | 0 | 2 | 3 | 0 | 5 | 0 | 1 | 1 | 0 | 1 | 35 | 0 | 0 | 10 | 2 |
|  | S<sup>b</sup> | 125 | 7 | 0 | 1 | 1 | 0 | 2 | 0 | 41 | 0 | 1 | 1 | 0 | 5 | 0 | 0 | 2 | 0 | 0 | 33 | 0 | 0 | 25 | 6 |
| Permethrin | R | 87 | 10 | 0 | 0 | 1 | 0 | 0 | 0 | 34 | 0 | 1 | 0 | 0 | 9 | 0 | 0 | 0 | 0 | 0 | 15 | 0 | 0 | 10 | 7 |
|  | S | 124 | 6 | 0 | 0 | 0 | 0 | 0 | 0 | 34 | 0 | 3 | 1 | 0 | 9 | 0 | 0 | 1 | 0 | 3 | 34 | 1 | 0 | 27 | 5 |
| Total |  | 465 | 37 | 0 | 2 | 4 | 0 | 2 | 0 | 161 | 0 | 7 | 5 | 0 | 28 | 0 | 1 | 4 | 0 | 4 | 117 | 1 | 0 | 72 | 20 |

<sup>a</sup>Resistant (alive), <sup>b</sup>Susceptible (dead), <sup>c</sup>Total number of mosquitoes. The order of the genotypes is shown for loci 1016/1534/410. Resistant allele at locus 1016 = I, 1534 = C, 410 = L, triple-resistant genotype II/CC/LL, triple-susceptible genotype VV/FF/VV.

## Discussion

In Colombia, the use of pyrethroids for the control of *A. aegypti* is a fairly recent phenomenon. Among the pyrethroids, λ-cyhalothrin and deltamethrin have most commonly been used to control *A. aegypti* in Colombia. However, resistance to λ-cyhalothrin has been more commonly reported than resistance to deltamethrin in Colombia, as demonstrated by results from previous studies [7, 9, 10, 11, 13, 18] as well as those obtained in the present study. In the findings presented here, we detected resistance to permethrin and λ-cyhalothrin in all populations and varying degrees of susceptibility to deltamethrin. This heterogeneity of resistance patterns within the pyrethroid class suggests that diverse mechanisms are contributing to these phenotypes.

Resistance to DDT is widespread in Colombia owing to the application of this organochlorine compound for more than five decades in the country [23]. DDT and pyrethroids share the mode of action consisting of delayed sodium channel closure and membrane repolarization [45]. The modification of this target site due to the presence of *kdr* mutations on the *para* gene can lead to cross-resistance to both DDT and pyrethroids. As such, the high prevalence of *kdr* alleles detected in our study may also be linked to previous selection pressures caused by DDT.

When our findings are compared with previous studies of insecticide resistance in *A. aegypti* in Colombia, our results are consistent with the findings of Maestre *et al.* [13] that reported resistance to λ-cyhalothrin in Barranquilla and Montería. However, those authors reported λ-cyhalothrin resistance and moderate resistance in Valledupar and Cartagena, respectively, whereas we detected resistance using the WHO bioassay but susceptibility using the CDC bioassay in both populations. Our results showing permethrin resistance were consistent with those reported by Maestre *et al*. [13] for Barranquilla and Montería; however, for Cartagena and Valledupar, Maestre *et al.* [13] reported susceptibility, whereas we observed resistance using the WHO bioassay but susceptibility using the CDC bioassay. For deltamethrin, Maestre *et al.* [13] reported resistance in Barranquilla; however, we found susceptibility using both bioassay methodologies. In Montería and Valledupar, Maestre *et al.* reported deltamethrin resistance in both populations, whereas we found susceptibility using the CDC bioassay and indications that resistance was developing using the WHO bioassay. In Cartagena, Maestre *et al.* [13] reported moderate deltamethrin resistance, whereas we observed resistance using the WHO bioassay and susceptibility using the CDC bioassay.

In Colombia, most previous insecticide susceptibility studies conducted on adult *A. aegypti* mosquitoes have used the CDC bioassay methodology, with the WHO bioassay methodology employed to a lesser degree. Typically, using both techniques, resistance to DDT has been observed in all *A. aegypti* populations evaluated in the country, together with variable susceptibility to pyrethroids and susceptibility to organophosphates in most populations [7].

In the present study, some discrepancies were observed between the results obtained with the WHO and CDC bioassay methodologies, indicating that the two techniques may not always provide consistent results. In studies by Aizoun *et al*. [42] and Fonseca *et al.* [23]., WHO and CDC bioassays were compared to determine the susceptibility of *Anopheles gambiae* to deltamethrin and *Anopheles nuñeztovari* to fenitrothion. Both studies reported susceptibility when using the WHO bioassay and resistance when using the CDC bioassay. The authors observed that the exposure time of the mosquitoes to the insecticide (diagnostic time) was considerably shorter in the case of the CDC bioassay, which could have led to an overestimation of resistance; although in fact the opposite was observed in our study. Despite the shorter exposure time in the CDC bioassay, populations that were classified as resistant in the WHO bioassay were classified as susceptible in the CDC bioassay. This could potentially be explained due to the mechanisms underlying the resistance; for example, resistance that is primarily caused by *kdr* would likely result in populations that are not quickly knocked down and thus scored as ‘resistant’ at 30 minutes. However, if the main mechanisms of resistance are metabolic, mosquitoes may initially be knocked down but could recover over time as their detoxification enzymes metabolize the insecticide. Indeed, our biochemical assay data suggest that elevated enzymatic activity is present in the populations that were studied.

Most previous studies regarding enzymatic activity have been conducted on *A. aegypti* populations from other regions of Colombia where alterations were detected, mainly in MFOs and nonspecific esterases, in populations from Antioquia, Chocó, Putumayo, Cauca, Valle del Cauca, Nariño, Huila, Santander, Meta, and Casanare [9–12]. The one previous study conducted in the Caribbean region of Colombia reported altered α-esterases and MFOs in *A. aegypti* from Valledupar, MFOs in Cienaga, and GSTs in Sincelejo. In Cartagena, Monteria, Barranquilla, San Juan, Puerto Colombia, and Soledad, no alterations in enzyme activity were detected [13]. Our results are consistent with the finding of highly altered MFOs in Valledupar, and we also detected altered β-esterases in that same population. We also detected highly altered α-esterases, β-esterases, MFOs and GSTs in Monteria; altered β-esterases and GSTs in Barranquilla; and altered GSTs in Cartagena. Additionally, in the present study we detected altered pNPA-esterases in the population of Juan de Acosta.

Regarding esterases, studies to date have reported the overexpression of β-esterases in populations resistant to organophosphates and pyrethroids [9–11]. Altered levels of α-esterase activity were detected previously in Valledupar in the study conducted by Maestre *et al.* [13]. In other countries, altered α-esterases, β-esterases, and MFOs have been reported in *A. aegypti* populations resistant to organophosphates, carbamates, and pyrethroids [39, 48–54].

There are no studies in Colombia incriminating insensitive acetylcholinesterase as a mechanism associated with resistance to organophosphates and carbamates in *A. aegypti.* A study by Grisales *et al.* [27] reported resistance to temephos in the population of *A. aegypti* from Cucuta (RR: 15X) without evidence of insensitive acetylcholinesterase, although they did detect esterase and oxidase-based mechanisms.

*Kdr* mutations are important mechanisms involved in DDT and pyrethroid resistance. In Colombia, the first *kdr* mutation reported in populations of *A. aegypti* was V1016I, which was identified in populations from Puerto Colombia, Soledad, Barranquilla, Valledupar, San Juan, Sincelejo, Montería, Cienaga and Cartagena, which are all located in the Caribbean region. In that initial report, the V1016I mutation showed frequencies ranging between 0.07 and 0.35; the lowest frequency was found in the Cienaga population and the highest was found in Soledad, Montería, and Barranquilla, with frequencies of 0.35, 0.33, and 0.32, respectively [13]. The highest frequency of 1016I that we detected in the present study was in Montería, with a frequency of 0.70, showing a large increase in the frequency in this population from what was originally reported by Maestre *et al.* [13]. In addition, an increase in the frequency of 1016I from 0.09 to 0.16 was detected in Cartagena and a reduced frequency was detected in Barranquilla and Valledupar, from 0.32 and 0.27, respectively, to 0.15 in both populations. V1016I had also previously been reported in Quindío at low levels of frequency (0.02–0.05) [29].

The F1534C mutation was first detected in Colombia in the department of Sincelejo (Sucre), in the Caribbean region [31]. It had also previously been reported in *A. aegypti* populations from Puerto Colombia, Soledad, Barranquilla, Valledupar, San Juan, Sincelejo, Montería, Cienaga and Cartagena with frequencies ranging between 0.74 and 0.88. When compared with the results reported previously, we observed increased frequencies of 1534C, having risen in Barranquilla from 0.74 to 0.76, in Cartagena from 0.86 to 0.97, in Montería from 0.88 to 1.00, and in Valledupar from 0.82 to 0.94. These increases are likely attributable to the constant pressure exerted by pyrethroid insecticides, which were heavily applied during the period between the two studies for the control of dengue, chikungunya, and Zika. Although there are no previous studies reporting this mutation in Juan de Acosta and Chiriguana, these populations also showed high frequencies (0.76 and 0.95, respectively). Moreover, high frequencies of 1534C have been reported in other areas of Colombia, including Villavicencio, Riohacha, and Bello, with frequencies of 0.63, 0.71, and 0.56, respectively [15]. In these latter three populations, the V410L mutation was also identified in Colombia for the first time, with frequencies of 0.46, 0.30, and 0.06, respectively. It is noteworthy that in that study,

*A. aegypti* from Bello were susceptible to λ-cyhalothrin, whereas those from Riohacha and Villavicencio were resistant. In these latter two populations, the researchers detected a positive association between V410L and V1016I and resistance to λ-cyhalothrin. In the present study, the V410L mutation was detected for the first time in the study populations, with frequencies ranging between 0.05 in Valledupar and 0.72 in Montería. The frequencies of the V1016I mutation were very similar to those of the V410L mutation in all the evaluated populations; this result is consistent with the findings reported by Granada *et al.* [15] for *A. aegypti* in Bello, Villavicencio, and Riohacha. Haddi *et al.* [38] reported the presence of the V410L mutation in resistant *A. aegypti* in Brazil and observed that this mutation, either alone or in combination with the F1534C mutation, was strongly associated with increased the resistance to type I and II pyrethroids. This is consistent with the results of the present study, where the 1534C and 410L alleles were associated with resistance to permethrin in the population of Juan de Acosta. The 1016I, 1534C, and 410L alleles were all associated with resistance to permethrin in the Chiriguana, Montería, and Valledupar populations based on phenotyping by the WHO bioassay. In addition, F1534C was associated with resistance to deltamethrin in Chiriguana, Valledupar, and Montería; V1016I and V410L were also associated with deltamethrin resistance in the case of the latter population. Similarly, an association was found between all three mutations and resistance to λ-cyhalothrin in Valledupar, Montería, and Juan de Acosta. This last result is consistent with the results of the study by Maestre *et al.* [55] which detected a significant positive correlation between the frequency of the 1016I allele and resistance to permethrin, λ-cyhalothrin, and cyfluthrin. However, no significant correlation was observed in that same study between 1534C and resistance to any pyrethroids [55].

Recent studies conducted in Mexico proposed three sequential models to explain the evolution of the V1016I, F1534C, and V410L mutations. The first model suggests that F1534C appeared first, providing low resistance levels, followed by the appearance of V1016I, which provided higher levels of resistance. The second model challenges the first model and proposes that V410L and V1016I occurred independently on a C1534 haplotype followed by cis conversion by recombination. Finally, a third model assumes that the three mutations appeared independently at low frequencies and that two recombination events rearranged them in a cis configuration [56]. Considering these previous models and the results obtained in the present investigation, it is possible to hypothesize that the appearance of V410L and V1016I did not occur independently because their allelic frequencies were so similar and they almost always appeared together.

Regarding the 1016/1534/410 phenotype–genotype association, a relationship between the VI_1016_/CC_1534_/VL_410_ genotype and resistance to λ-cyhalothrin and permethrin was detected in the present study. These results are consistent with the study conducted by Haddi *et al.* [38] in a pyrethroid-resistant *A. aegypti* strain from Brazil, where V410L alone or in combination with F1534C was shown to reduce sodium channel sensitivity to type I (permethrin) and type II pyrethroids (λ- cyhalothrin and deltamethrin). In addition, these results further support the notion that the presence of VI_1016_ and VL_410_ heterozygotes is sufficient to confer resistance to deltamethrin [56]. These findings suggest that the interactions of multiple mutations play a role in the response of *A. aegypti* sodium channels to insecticides [57].

## Conclusions

Variability was observed in pyrethroid susceptibility using the WHO and CDC bioassay methodologies, highlighting the importance of using a consistent methodology to routinely screen populations for susceptibility. The altered activity levels of β-esterases, α-esterases, MFOs, and GSTs suggest that metabolic resistance may be important in these populations. The *kdr* mutations V1016I, F1534C, and V410L were detected in all populations, with 1534C being nearly fixed in all except two populations. Finally, associations were observed between the F1534C mutation and resistance to permethrin in all populations, the F1534C mutation with resistance to deltamethrin in Chiriguana, Montería, and Valledupar, and the V1016I, F1534C, and V410L mutations and resistance to λ- cyhalothrin in Juan de Acosta, Valledupar, and Montería.

## Acknowledgments

The authors thank Patrica Fuya from the Laboratory of Entomology, Colombian National Institute of Health; Jhon Jairo González from the Caribbean Region Node, of the project “Strengthening the entomological surveillance of *Aedes aegypti* in Colombia for the reinforcement of the national entomology network”; Lucrecia Vizcaino of the Entomology Branch of the U.S. Centers for Disease Control and Prevention; the Public Health Laboratory group, Entomology Laboratory of the District of Barranquilla and the Department of Atlántico; and Sergio Goenaga of the Public Health Laboratory of Atlántico.

## Author Contributions

LSV and PXPL conceived and designed the study; LSV and AL obtained financial support; PXPL performed the fieldwork; PXPL and GRV performed the laboratory work; and PXPL and RMS analyzed the data and its presentation. PXPL, DGC, RYMS, and AL drafted the manuscript. All the authors have provided critical information on the findings and have read and approved the final manuscript.

## Disclaimer

The findings and conclusions in this paper are those of the authors and do not necessarily represent the official position of the Centers for Disease Control and Prevention.

